# Sky islands of Southwest China. II: Unraveling hidden species diversity of talpid moles using phylogenomics and skull-based deep learning

**DOI:** 10.1101/2025.03.06.641773

**Authors:** Kai He, Anlong Li, Quentin Martinez, Xiaoyun Wang, Zhongzheng Chen, Shuiwang He, Sining Xie, Zeling Zeng, Kunhui Wang, Ziqi Ye, Hao Ruan, Shiyun Liu, Qiuqin Lu, Xiaoyun Zheng, Jiayi Luo, Wenyu Song, Achim Schwermann, Haibin Yu, Wenhua Yu, Mark Springer, Shaoying Liu, Song Li, Feiyun Tu, Zhong Cao, Kevin L. Campbell

## Abstract

The sky islands of Southwest China, characterized by dramatic topographical and climatic variations, are prominent hotspots of biodiversity and endemism. Organisms inhabiting middle-to-high elevation habitats in this region are geographically isolated within distinct mountain chains, which over geological time have been subjected to isolation-by-distance and isolation-by-environment. These processes have led to profound allopatric diversification and strong phylogeographic structuring, resulting in a plethora of genetically distinct cryptic species, as is becoming increasingly evident for many small mammal families. However, morphological conservatism can pose significant challenges in delineating these clades within species complexes. In this study, we leverage artificial intelligence technologies to unravel the hidden species diversity of moles (family Talpidae) in Southwest China’s sky islands. We first employed ultraconserved elements (UCEs) to investigate the evolutionary history of talpid moles, conducted molecular species delimitation using mitochondrial and multi-locus genes, and utilized both traditional and geometric morphometrics to examine their morphological disparity. To address the challenges of morphology based cryptic species identification, we developed a deep learning Hierarchical Identification of Species NETwork (HIS-NET) to create an image-based model that analyzes four different views of the skull/mandible to distinguish genera and species hierarchically. HIS-NET not only achieved expert-level accuracy in species identification but also effectively distinguished between cryptic and known species, aiding in the identification of key morphological variation intervals. Our results support the recognition of allopatrically distributed taxa in *Euroscaptor* and *Parascaptor* as full species, thereby confirming that species diversity in this region remains underestimated. Beyond advancing our understanding of speciation in this unique and fragile ecosystem, our study serves as a proof-of-concept, demonstrating the power of deep learning in unraveling hidden biodiversity within this and other species complexes.

## 1 Introduction

The mountainous regions of Southwest China, situated adjacent to and east of the Qinghai-Tibet Plateau, are characterized by a complex geomorphological history that produced dramatic topographical landscapes ranging from 300 m to >7,000 m above sea level ^[1]^. Shaped by the uplift of the Qinghai-Tibet Plateau, this region comprises a series of mountain chains that are physically isolated from one another by basins, valleys and extensive drainage systems that predominantly run in north-south directions ^[2]^. This uneven terrain supports varied climates across both vertical and horizontal space, providing a mosaic of ecologically diverse habitats ^[3]^. Another distinctive feature of this region is the discontinuous distribution of similar habitats at medium to high elevations across different mountain areas, creating a unique geographical backdrop that has earned it the "Southwest China sky-island complex" designation ^[4]^.

The concept of sky islands, originally coined to describe isolated mountain ranges in the southwestern United States and northern Mexico ^[5, 6]^, applies particularly well to these mountainous regions of Southwest China. Indeed, the progressive uplift of this region through geological time created an archipelago-like array of high-elevation “island” ecosystems separated by intervening lowlands that led to the fragmentation of plant and animal populations inhabiting middle-to-high elevation environments across the mountain ranges ^[7]^. As such, this geographic panorama, in conjunction with climatic fluctuations such as the Pleistocene glaciations, has significantly influenced evolutionary processes and speciation events, resulting in a complex tapestry of endemic and relict species across these sky islands ^[8, 9]^.

The Southwest China sky island complex has spawned globally important biodiversity hotspots ^[10]^ that harbor an exceptionally high level of species diversity. For example, this region is home to over 12,000 described species of vascular plants, of which 29% are endemic ^[11]^, and hosts approximately 50% of China’s bird and mammal species ^[12]^. However, the full extent of biodiversity in this region remains underexplored, largely due to the presence of cryptic species ^[13]^ – morphologically similar but genetically distinct lineages – which pose a significant challenge to biodiversity assessments and conservation efforts ^[14, 15]^.

The primary mechanisms driving diversification and speciation in sky island complexes are isolation-by-distance (IBD) and isolation-by-environment (IBE), describing isolation driven by geographic distance or as a product of environmental dissimilarity, respectively ^[16]^. Both processes may eventually result in allopatric speciation as populations of organisms in the newly isolated habitats can diverge into distinct genetic lineages across the chain of sky-islands. This evolutionary pattern is evident in many plants, insects, amphibians, and small mammals inhabiting the mountainous region of southwestern China ^[17–21]^. Accordingly, previously recognized species with broad distributions are now classified into a complex of multiple species that exhibit patterns of allopatric speciation across different mountain islands ^[22]^. For instance, the rodent genus *Typhlomys*, originally thought to comprise a single species widely distributed throughout southern China, has been found to comprise at least seven distinct species and undescribed cryptic lineages ^[23, 24]^. All of these newly recognized species show some degree of geographical isolation, with each genetic lineage confined to a specific mountainous area. These findings imply that other widely distributed small mammal families inhabiting this region may also house undescribed species, thereby hampering our ability to fully assess the species diversity in the Southwest China sky island complex. The mole family Talpidae serves as a prime example, as cryptic diversity is likely widespread in this region but remains largely unresolved ^[8, 14, 15]^.

Classified within the order Eulipotyphla, talpids are widely distributed across Europe, Asia, and North America and are characterized by a range of ecological and morphological specializations. These include the shrew-like moles of the monogeneric (*Uropsilus*) subfamily Uropsilinae which possess echolocation abilities ^[25]^, a unique trait in the family, though are primarily terrestrial in their ecological niche. The remaining clades belong to the subfamily Talpinae and have evolved to exploit a diverse variety of habitats: the semi-fossorial shrew moles (tribes Scaptonychini, Urotrichini, and Neurotrichini), the semi-aquatic desmans (Desmanini), and the fully fossorial (Talpini and Scalopini) and semi-aquatic/semi-fossorial (Condylurini) moles.

As of 2018, Wilson and Mittermeier ^[26]^ recognized 54 extant talpid species within 18 genera globally (**Table S1**) though other recent studies have revealed additional new ^[27–32]^ and cryptic species ^[8, 14, 15, 33–35]^ and even a new genus (*Alpiscaptulus*)^[36]^. The mountains of Southwest China, which harbor 17 currently recognized species, are not only a hotspot of this family’s species diversity but are also home to the highest number of cryptic species, particularly within *Uropsilus* ^[14, 33]^, *Euroscaptor* ^[37]^, *Parascaptor* ^[8, 34, 35]^, and *Scaptonyx* ^[8, 15]^. While a number of distinct talpid lineages identified in these studies have since been recognized as full species – e.g., *Uropsilus nivatus*, *U*. *atronates* ^[26]^, *Euroscaptor orlovi*, *E. kuznetsovi* ^[38]^*, Scaptonyx wangi* and *S. affinis* ^[27]^ – others are yet to be investigated. Importantly, species in these genera are middle-to-high elevation inhabitants that exhibit clear sky-island distribution patterns ^[14, 15]^. This discontinuous distribution results in diversification following basic IBD or IBE principles, that has led to strong genetic structuring across the region. However, the similar habitats found among the many montane islands have also fostered morphological conservatism ^[39]^, resulting in many of these same genetically distinct lineages having a high degree of similarity in external and craniodental features. Thus, while molecular-only based approaches have detected cryptic species following the phylogenetic species concept (PSC), they often leave open the question of whether these genetic lineages are morphologically diagnosable. This challenge is particularly pertinent when considering either the diagnostic morphological phylogenetic species concept (dmPSC ^[40]^) or the unified species concept (USC ^[41]^), which both require diagnosable morphological differences among putative species, thereby underscoring the need for tools that can bridge the gap between molecular structuring and full species validation.

In recent years, deep learning has revolutionized classification and identification across various fields ^[42]^. Convolutional neural networks (CNNs), in particular, have demonstrated remarkable efficacy in image classification tasks, even without additional contextual information ^[43, 44]^. The power of CNNs lies in their ability to automatically learn hierarchical feature representations from raw image data ^[45]^, making them particularly well-suited for tackling complex pattern recognition problems in biology, including species identification. Indeed, CNNs have been successfully applied to the identification of plant ^[46]^ and animal species ^[47]^, demonstrating their efficacy and broad potential in species identification. However, despite their successful use in invertebrates ^[48]^, the application of CNNs in the identification of vertebrate species remains surprisingly limited. Only a few studies have explored the potential use of CNNs for the identification of mammal and reptile species, and these studies typically included only a small number of closely related or congeneric species ^[49, 50]^. Accordingly, there is a conspicuous lack of comprehensive, family-level studies that explore the capacity of CNNs to discern morphological differences at both species and higher taxonomic levels.

The classification system in taxonomy is inherently hierarchical, progressing from species to genus to family levels and beyond. Taxonomists recognize that morphological differences at various taxonomic levels, such as between genera and between congeneric species, are distinctly different. Hierarchical classification using deep learning for multi-label image categorization is a well-established approach in the information science domain, and can significantly enhance information retrieval efficiency and accuracy ^[51]^. However, the application of hierarchical classification strategies to species identification has been surprisingly limited with only a few studies having explored this approach in the context of biological taxonomy ^[52, 53]^. Additionally, most previous studies have predominantly relied on either single or multiple photographs of the same view, potentially missing crucial diagnostic features that are only evident from different angles or on structures obstructed from view. To our knowledge, few studies have examined whether integrating multiple views of specimens capture more key diagnostic characters and improve identification accuracy ^[54]^.

To address these limitations and explore the full potential of CNNs in vertebrate taxonomy, we conducted an integrative evolutionary history study on the Talpidae family with a focus on exploring the cryptic species diversity in the Southwest China sky island complex. We first generated an ultraconserved element (UCE) dataset to investigate the molecular phylogenetics of talpids and applied species delimitation methods employing mitochondrial and nuclear genes to confirm any genetically distinct lineages observed (**Table S2**). We then applied both traditional and geometric morphometrics to test whether the identified cryptic species are also phenotypically distinct from congenerical recognized species. Our final aim was to evaluate the potential of deep learning techniques to uncover morphological differences that may be imperceptible to traditional methods to accurately distinguish previously recognized from cryptic talpid species. To this end, we developed a novel **H**ierarchical **I**dentification of **S**pecies **NET**work (**HIS-NET**) model using CNN, which integrates multiple views of the skull and mandible of voucher specimens for species identification. This research not only advances our understanding of talpid diversity in the Southwest China sky island complex but also provides a proof-of-principle for the application of deep learning in resolving complex taxonomic challenges.

## Material and Methods

### 2.1 Molecular phylogeny and species delimitation

#### 2.1.1 Taxonomy and taxon sampling

We followed the taxonomy proposed by Wilson and Mittermeier ^[26]^ but included updates in subsequent studies thereafter, expanding the Talpidae family to 66 species within 19 genera **(Table S1**). We propose several candidate species within *Euroscaptor* (n=2), *Parascaptor* (n=2), and *Uropsilus* (n=1) in southwestern China, based on their genetic and, where applicable, morphological distinctiveness. We collected a total of 61 specimens representing 60 species/candidate species for high-throughput sequencing (**Table S2**), that included 44 talpid species, 11 shrews (Soricidae), 5 erinaceids (Erinaceidae), and 1 solenodon (Solenodontidae). Tissue samples preserved in 95% ethanol were obtained from various sources, including loans from the following institutions: Kunming Institute of Zoology (KIZ, China), National Museum of Nature and Science (NMNS, Japan), National Museum of Natural History (USNM, USA), Burke Museum of Natural History and Culture (NWBM, USA), Field Museum of Natural History (FMNH, USA), New Mexico Museum of Natural History (NMMNH, USA), as well as several personal collections (**Table S2**). To test the validity of cryptic species, we also collected specimens from southern China and Myanmar.

#### 2.1.2 DNA Library preparation

Total genomic DNA was extracted from tissue samples using a Qiagen DNeasy Blood and Tissue Kit (Qiagen). The extracted DNA was randomly fragmented to sizes ranging from 200 to 400 bp using the NEBNext dsDNA Fragmentase (New England Biolabs, Canada). Library construction was performed using the NEBNext Fast DNA Fragmentation & Library Prep Set for Ion Torrent (New England Biolabs, Canada), with each library incorporating a unique barcode adapter from the NEXTflex DNA Barcodes for Ion Torrent (BIOO Scientific, USA). We conducted size selection using a 2% E-gel on an E-Gel Electrophoresis System (Invitrogen, Canada). We re-amplified the libraries using a NEBNext High-Fidelity 2X PCR Master Mix (New England Biolabs, Canada), and purified using Dynabeads Magnetic Beads. Following library construction, we determined the concentrations using a Qubit 4 fluorometer and Qubit dsDNA Assay (Thermo Fisher Scientific, Canada).

#### 2.1.3 UCE capture and sequencing

For ultra-conserved elements (UCE) capture, we utilized myBaits probes synthesized by Arbor Biosciences (Ann Arbor, MI, USA). Four DNA libraries of similar quantity were pooled prior to hybridization. In-solution hybridization was performed following the myBaits manufacturer’s protocol. The enriched libraries were purified using Dynabeads Magnetic Beads and amplified with a NEBNext High-Fidelity 2X PCR Master Mix (New England Biolabs, Canada). The indexed captured libraries were pooled in equal concentrations and sequenced on an Ion Torrent PGM or Ion Torrent Proton sequencer.

#### 2.1.4 Data processing and phylogenetic analysis

Raw data were automatically de-multiplexed and converted to FASTQ format on the Torrent Suite v4.0.2 software (Thermo Fisher Scientific, Canada). Given that Ion Torrent platforms produce single-end (rather than pair-end) reads, we pre-processed the data using packages optimized for single-end reads following He et al. ^[37]^. Initially, we trimmed adapters and barcodes using AlienTrimmer v0.4 ^[55]^ as part of the ClinQC v.1 package with conservative parameters (-k 15 -m 5 -l 15) ^[56]^. We then removed poor quality data, using the DynamicTrim function of SolexaQA + +v3.1 ^[57]^, and eliminated duplicates using ParDRe ^[58]^. Sequence correction was performed using karect ^[59]^ and the data were used for downstream assembly.

For the UCE assembly, we followed the PHYLUCE v1.7 ^[60]^ protocol "UCE Phylogenomics" to extract UCE loci. Data were de novo assembled using SPAdes v3.1 ^[61]^. The original “uce-5k-probes” were used as references to extract UCE loci from the draft assembled contigs. Additionally, UCE loci were extracted from publicly available (GenBank) genomes of *Condylura cristata*, *Sorex araneus*, *Erinaceus europaeus*, and *Solenodon paradoxus*, following the PHYLUCE tutorial "Harvesting UCE loci from genomes". Loci present in at least 50% of taxa were retained for further analyses. Each locus was trimmed using ClipKIT with the “kpic-smart-gap” parameter to retain parsimony-informative sites with few gaps ^[62]^. Loci were then realigned using CIAlign to remove divergent sequences, insertions, and sequences shorter than 100 bp ^[63]^. After data processing, 2989 UCE loci were retained, with an average of 2049 UCE loci per sample.

#### 2.1.5 Phylogenetic analyses

We conducted both concatenation and summary-coalescent species tree analyses using a two-step strategy. First, we estimated gene trees for each UCE locus using RAxML-NG with the following parameters: --brlen scaled --bs-trees autoMRE(1000) --bs-metric fbp,tbe ^[64]^. This included simultaneous rapid bootstrap analyses and searches for the best scoring maximum likelihood tree (--all). Subsequently, we employed TreeShrink ^[65]^ to remove taxa represented by very long branches from the corresponding UCE locus alignments. The pruned alignments were then used for tree estimations.

For concatenated species tree estimation, we divided each UCE into three data blocks (core, left flanking, and right flanking regions) using SWSC-EN ^[66]^. We determined the best partitioning scheme using PartitionFinder 2 ^[67]^ with the rclusterf search under the GTR+G model based on AICc. The optimal scheme included 1032 partitions. The concatenated tree was then estimated using RAxML-NG as described above.

For coalescent species tree estimation, gene trees for each UCE locus were estimated with RAxML-NG. Branches with bootstrap supports lower than 10 were collapsed using nw_ed ^[68]^. The species tree was then estimated using ASTRAL-IV v1.19 ^[69]^.

#### 2.1.6 Molecular species delimitation

To extend our taxon sampling, we included sequence data from newly collected specimens captured in Yunnan, Sichuan, and Myanmar representing *Parascaptor* sp.1, *P*. sp.2 and *Euroscaptor* sp.1 in our species delimitation analyses (**Table S2**). For each specimen, we amplified the complete mitochondrial *CYTB* gene and also amplified 11 nuclear genes (*ADORA3*, *APP*, *ATP7A*, *BCHE*, *BDNF*, *BMI1*, *BRCA1*, *CREM*, *PLCB4*, *RAG1* and *RAG2*) for a subset of samples using the primers presented in ^[8, 70]^. We were not able to collect additional *Euroscaptor* sp. 2 specimens for this analysis.

We also retrieved available *CYTB* sequences for *Euroscaptor* (n=86) and *Parascaptor* (n=10) from GenBank and combined them with our newly generated sequences for phylogenetic analysis (**Table S2**). We first aligned the sequences using MUSCLE ^[71]^, and estimated the gene trees using RAxML as described above. We then employed two single-locus species delimitation methods: Automatic Barcode Gap Discovery (ABGD; ^[72]^) and Assemble Species by Automatic Partitioning (ASAP; ^[73]^). Additionally, we calculated pairwise K2P distances and generated a genetic distance heatmap in R v4.4 using the packages ape ^[74]^ and ggplot2 ^[75]^. The *Parascaptor* and *Euroscaptor* datasets were analyzed separately.

Finally, we performed coalescent-based species delimitation analyses using 11 nuclear genes, as implemented in BPP v4.7 ^[76]^. Sample sizes for *Parascaptor* sp.1 and *Parascaptor* sp.2 were 4 and 16, respectively. The *Euroscaptor* analysis included 22 specimens: *E. malayana* (n=2), *E. klossi* (n=1), *E. kuznetsovi* (n=8), *E. orlovi* (n=1), *E. longirostris* (n=4), and *Euroscaptor* sp.1 (n=6). We did not include *Euroscaptor* sp.2 because we had only two samples in both the morphological and molecular data sets thereby precluding a robust examination of its taxonomic status in this study. For the *Parascaptor* dataset, we employed the A10 model, which focuses solely on species delimitation since only two putative species were included. For the *Euroscaptor* dataset, we utilized both the A10 and A11 models, with the latter allowing for simultaneous species delimitation and species-tree estimation. We applied both algorithms 0 and 1, and conducted six analyses for each dataset per model, using various combinations of parameters and priors (**Table S3**).

### 2.2 Morphometric and geometric morphometric analyses

#### 2.2.1 Traditional morphometrics

Fifteen craniomandibular variables were measured using a digital caliper graduated to 0.01 mm from 66 specimens of *Parascaptor* and 76 specimens of *Euroscaptor* (Appendix I): CIL (Condyloincisive length), PIL (Palatoincisive length), PPL (Postpalatal length), CB (Cranial breadth), IOB (Interorbital breadth), ZB (Zygomatic breadth), CH (Cranial height), UTL (Upper toothrow length), P4-M3 (Distance from the upper fourth premolar to the upper third molar), M2M2 (Maximum width across the upper second molars), BFM (Foramen magnum breadth), LTR (Lower toothrow length not including first incisor), LLM (Lower molars length), ML (Mandible length), HCP (Height of coronoid process). We conducted Principal Component Analysis (PCA) and Canonical Variate Analysis (CVA) on log10-transformed variables using the stats and Morpho packages ^[77]^, respectively. The results were visualized using ggplot2 and plotly.

#### 2.2.2 Specimen accession and photography

We photographed specimens housed at natural history museums in China, Vietnam, Japan, Germany, and the USA (Appendix I). Specimens at the Institute of Zoology, Chinese Academy of Sciences (IOS); Kunming Institute of Zoology, Chinese Academy of Sciences (KIZ); Sichuan Academy of Forestry (SAF); Guangdong Institute of Zoology (GIZ); National Museum of Natural Science at Taichung (NMNST); Smithsonian Institution National Museum of Natural History (USNM); American Museum of Natural History (AMNH); Field Museum of Natural History (FMNH); Museum of Comparative Zoology, Harvard (MCZ); National Museum of Nature and Science of Japan (NSMT); Hokkaido University Natural History Museum (HUNHM); University of Miyazaki; Institute of Ecology and Biological Resources of Vietnam (IEBR) were photographed by K.H. Photos at the State Museum of Natural History Stuttgart (SMNS) were taken by Q.M. Additional photos of two Russian desman specimens were taken by T. Martin and C. Steinweg, while photos of two *Galemys* specimens were taken by J. Decher and C. Montermann.

We utilized either a Nikon D300 or D7100 with a Nikon 105mm f/2.8G lens, or a Canon EOS 7D Mark II with an EF 100mm f/2.8L IS USM. Consistent and standardized criteria were applied for dorsal, ventral, and lateral views of the skull, as well as the lateral view of the mandible. A camera stand was used to stabilize the camera, with the skull or mandible placed on a small plate approximately 10-30 cm below the lens, depending on the specimen size. All pictures were taken on a blue background. A level was placed on both the camera and the specimen plate to ensure the lens was perpendicular to the specimens. Additional lighting or a flash was used when appropriate. Depending on the lighting and flash systems, which varied in different museums, we adjusted the F-stop in the range of 13 to 23 and ISO from 200 to 1250 to ensure high photo quality. Due to the requirements of deep learning, we included only species with at least three specimens, excluding *Alpiscaptulus medogensis* and *Euroscaptor* sp.2, which each had only two specimens available. In total, we collected 2998 photos representing 819 specimens from 18 (out of 19) talpid genera and 51 species/putative species.

#### 2.2.3 Geometric morphometric analysis

We used the ventral view of the skulls for geometric morphometric analyses. Only complete skulls were included for *Parascaptor* and *Euroscaptor*. A total of 38 specimens representing *P. leucura* (n=13), *Parascaptor* sp.1 (3), and *Parascaptor* sp.2 (22), and 55 specimens representing *E. grandis* (5), *E. klossi* (8), *E. kuznetsovi* (3), *E. longirostris* (12), *E. malayana* (6), *E. micrura* (10), *E. orlovi* (3), *E. parvidens* (4), and *Euroscaptor* sp.1 (4) were used. The backgrounds of the images were removed using Adobe Photoshop. The analyses were conducted as implemented in R v4.4. We utilized the outlineR package to generate the outlines of the skulls. Subsequently, the Momocs package was employed to subsample 150 coordinates from the existing points per specimen ^[78]^. We then performed Elliptical Fourier Analysis (EFA) to quantify the shape of each outline. Following the EFA, we conducted PCA and CVA as described above.

### 2.3 CNN based species identification

#### 2.3.1 Data labeling and manipulation

Each photo (see section 4.2.2) was labeled with the following information: genus and species name, voucher ID, skull or mandible, and the view of the skull (lateral, ventral, or dorsal). It is common for small mammal skull specimens in museums to be broken or have missing parts, as many were captured using snap traps. Among the 819 specimens included in the data set, 133 had fewer than four images due to missing or completely broken cranial bones, while 203 images were labeled as partly broken (**Table S5**). Minor damage such as missing zygomatic arches were not considered as broken. We manually cropped the images to retain only the skull or mandible, removing the peripheral areas. Then we standardized the images to square dimensions by padding the shorter edges with black pixels. The dataset was then divided into training (data Trn0) and testing (Tst0) sets with a ratio of 8:2. This division was performed at the specimen level, ensuring that all images of a specimen were placed into the same dataset (**Table S6**).

#### 2.3.2 Data augmentation

It is well-established that larger datasets lead to improved performance in deep learning networks ^[79]^. However, obtaining specimens of rare and newly recognized species, often represented by only a few samples in museum collections, presents a significant challenge. To address this limitation, we employed a comprehensive data augmentation strategy, introducing minor distortions to images of the training data to mitigate overfitting during neural network training ^[80]^. To achieve numerical balance among different genera, we applied higher data augmentation factors to those with fewer samples. Specifically, we utilized between 3 to 14 augmentation techniques per genus including rotation (rotate 90, rotate 45), mirroring (horizontal, vertical), masking (drop out, coarse drop out), blurring (gaussian, average, motion), noise injection (salt and pepper, additive gaussian, impulse), and contrast change (gamma contrast, sigmoid contrast) (**Fig. S12**). For genera with particularly few specimens (e.g., *Desmana*, *Galemys*, and *Scapanulus*), we created duplicates of the augmented data and applied an additional augmentation method, average pooling, to these duplicates, resulting in a 30-fold increase in the number of augmented samples. The augmentation factors ranged from 4 to 30 at the genus level, with an average augmentation factor of approximately 9 (**Table S5**).

#### 2.3.3 CNN selection

Different types of CNN architectures have varying feature extraction capabilities, with shallow CNNs better suited for capturing low-level features, while deeper CNNs excel at identifying more complex, high-level semantic information ^[81]^. To select the most suitable CNN for talpid mole species identification, we compared various networks with different architectures, complexities, and performance metrics. Specifically, we evaluated AlexNet, the EfficientNet series ^[82]^, GoogleNet ^[83]^, MobileNet ^[84]^, the ResNet series ^[85]^, the ShuffleNet series ^[86]^, and the VGGNet series ^[87]^. The parameters used per model are given in **Table S7**. To optimize computational efficiency, we first standardized the image resolution to 224x224 pixels for both the training and testing datasets. To improve network performance and species recognition accuracy, we utilized transfer learning ^[88]^, leveraging network parameters pre-trained on large datasets like ImageNet ^[89]^. Each network was trained and evaluated based on accuracy and computational efficiency. Additionally, we assessed the impact of data augmentation by using training datasets with (Trn1-224) and without (Trn0-224) data augmentation.

#### 2.3.4 Image resolution selection

Image resolution significantly impacts the performance of deep learning models ^[90]^. Higher image resolution offers richer details, potentially enhancing classification accuracy, but also increases computational complexity due to the larger number of model parameters. To investigate this trade-off, we compared network performance across different resolutions: 224x224, 260x260, 300x300, and 380x380 pixels (**Table S8**). Given that the EfficientNet series demonstrated the highest accuracy (see Results section), our analyses focused on EfficientNet B0, B2, B3, and B4. Briefly, these networks differ in their network depth, width, and resolution, with higher versions (B2, B3, B4) having increased capacity and complexity to capture more intricate features compared to B0. Since data augmentation significantly improved network accuracy (see Results section), we exclusively used augmented datasets in this and subsequent analyses.

#### 2.3.5 Individual reconciliation

The EfficientNet, as well as other CNNs, performs identification on an image-based manner, yielding probabilistic outputs. Given that each specimen is represented by up to four images—dorsal, ventral, and lateral views of the skull, and a ventral view of the mandible—and recognizing that each cranial and mandibular view contains varying quantities of diagnostically relevant characters for species identification, thus yielding differential identification accuracies, we developed a weighted approach to reconcile multiple image outputs into a consolidated specimen identification.

We calculated genus-level identification accuracy for each view and derived weights using the formula: *ω_i_* = 1 − (*Acc_max_* – *Acc_i_*) *η, where ω_i_ represents the weight for each view, *Acc_max_* denotes the highest accuracy among the four views, *Acc_i_* is the accuracy for each individual view, and η is a parameter used calibrate the weighting (set to 10 in this study). These weights (ω1, ω2, ω3, ω4) are then used to combine the probability distributions of four different views of the skulls. The final genus-level identification for each specimen is determined by maximizing the weighted sum of these views’ probability distributions. The process is expressed mathematically as follows: *P_individual_* = *argmax*{*P_s_d_* * ω_1_ + *P_s_v_* * ω_2_ + *P_s_l_* * ω_3_ + *P_m_l_* * ω_4_}, where *P_s_d_*, *P_s_v_*, *P_s_l_*, *P_m_l_* represent the probability distribution matrices for the dorsal, ventral, lateral skull, and lateral mandible views, respectively, and ω_1-4_ are the corresponding weights for each view as calculated earlier,. The sum of the probabilities in each matrix is normalized to 1. The ***argmax*** function identifies the class with the highest combined probability, which corresponds to the final predicted genus-level category for the specimen, denoted as *P_individual_*.

#### 2.3.6 A hierarchical identification network

Given that direct species-level identification resulted in an overall accuracy below 90% (see Results Section), and considering that seven of the 18 talpid genera are polytypic (*Euroscaptor, Mogera, Talpa, Scapanus, Scaptonyx*, and *Uropsilus* while *Parascaptor* was shown to contain two cryptic species), we implemented a hierarchical classification scheme, employing a cascade of classifiers to sequentially identify specimens to the genus and species level. This approach utilized two primary classifiers: one trained to distinguish among all talpid genera, and a second suite of species-level classifiers tailored to a specific polytypic genus. The genus classifier first assigned each specimen’s photographic set to the most probable genus. For specimens predicted to belong to a polytypic genus, the photographic set was then passed to the corresponding species-level classifier. Individual image classifications were then reconciled to produce a single, consolidated species prediction for each specimen. This strategy addresses the hierarchical nature of taxonomic relationships and the varying levels of morphological differentiation between and within genera. We named our network, **H**ierarchical **I**dentification of **S**pecies **NET**work (**HIS-NET**).

To assess the consistency of the network performance and minimize the influence of data partitioning on the result, we implemented a five-fold cross-validation approach. The dataset was partitioned into five subsets of approximately equal size. In each iteration, four subsets were utilized for network training, while the remaining subset served as the validation set.

To elucidate the decision-making process of our network, we employed the Class Activation Mapping (CAM) algorithm to generate heatmaps, which were superimposed on the original images. We then manually inspected these heatmaps to discern which part of the skull/mandible the deep learning network prioritized and identified as crucial for species identification.

## 3. Results

### 3.1 Talpid phylogenetic hypotheses employing UCE data are robust and support the existence of cryptic species

Both concatenation (RAxML; **Fig. 1**, **Fig. S1**) and coalescent (ASTRAL; **Fig. S2**) analyses produced highly supported topologies that were broadly consistent with each other. Briefly, both analyses recovered the subfamilies Uropsilinae and Talpinae, unite the three shrew mole tribes into a monophyletic clade, and support a basal divergence of Scalopini within Talpinae. Of note, *Condylura* was placed sister to shrew moles with high support in both analyses (maximum likelihood bootstrap value [BS]=100, ASTRAL coalescent bootstrap value [c-BS]=1.0). The only topological discrepancies between analyses pertain to shrew mole interrelationships and the placement of a single lineage (*Euroscaptor* sp.2) within Talpini. Chief among these is that the UCE concatenation analysis strongly supported Chinese shrew moles (*Scaptonyx*) as sister to North American shrew moles (*Neurotrichus*), while the UCE coalescent analysis placed *Scaptonyx* sister to the Japanese shrew moles (*Urotrichus* + *Dymecodon*) with high support. Importantly, both analyses revealed the presence of distinctive genetic lineages in *Uropsilus*, *Euroscaptor* and *Parascaptor*, confirming an underestimation of species diversity within the Southwest China sky island complex. Given that the *Uropsilus* species continuum comprises numerous undescribed species ^[14, 32]^ that were not sampled here, we will henceforth focus our attention on cryptic species delimitation within *Euroscaptor* and *Parascaptor*.

**Figure 1.**
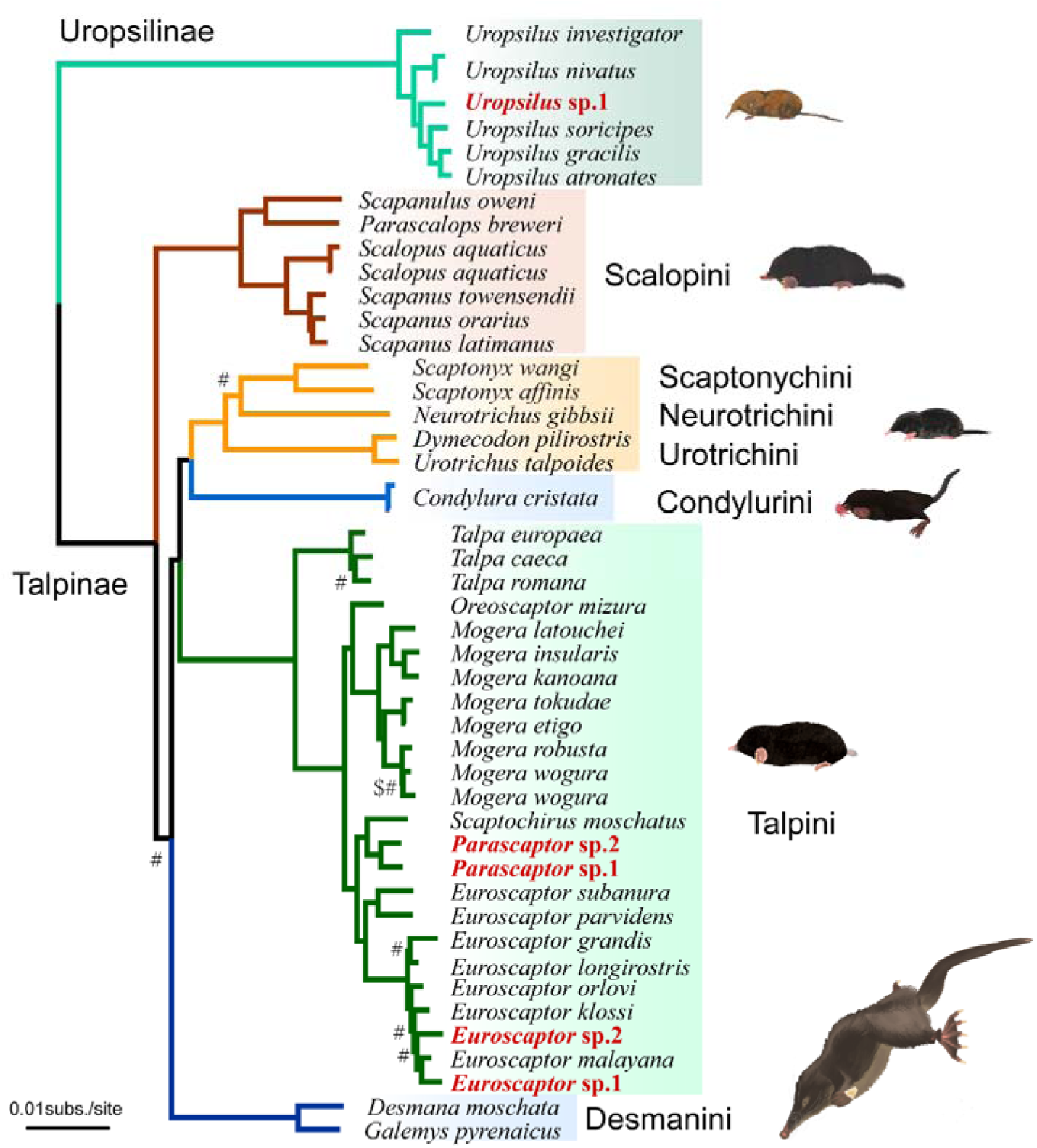
Phylogenetic relationships among talpid species. This maximum likelihood (RAxML) tree was constructed using concatenated ultraconserved element (UCE) data. Unless specified, all relationships are highly supported by both concatenation and coalescent analyses. Relationships not strongly supported in concatenation (BS <0.95: $; Fig. S1) or coalescent analysis (c-BS <0.95: #; Fig. S2) are indicated. Paintings of representative species by Umi Matsushita.

### 3.2 Molecular species delimitation supports the genetic distinctiveness of cryptic species

To test whether the genetically distinctive cryptic lineages identified in *Euroscaptor* and *Parascaptor* form monophyletic clades according to the PSC, we extended our taxon sampling for members of these genera and estimated mitochondrial phylogenies based on cytochrome B (*CYTB*) gene sequences. The *CYTB* gene tree revealed that the *Parascaptor* sp.1 clade represents a basal divergence of the genus (**Fig. S3A**) that is comprised of specimens from eastern Yunnan and southern Sichuan (**Fig. S4A**). *P. leucura* is represented by one specimen from northeast India that was placed sister to *P.* sp. 2 specimens from southwestern Sichuan, central and western Yunnan, and northeastern Myanmar (**Fig. S4A**). Notably, both species delimitation analyses – Automatic Barcode Gap Discovery (ABGD) and Assemble Species by Automatic Partitioning (ASAP) – and our K2P genetic distance heatmaps further revealed distinctive subclusters among *Parascaptor* sp.2 populations from Yunnan, Sichuan, and Myanmar (**Fig. S3A**). However, due to limited specimen sampling from most identified subclusters, we here conservatively treat *Parascaptor* sp.2 as a single putative species.

The *Euroscaptor* sp.1 and *E.* sp.2 samples used in the UCE analyses are from museum specimens collected in the 1960’s, with *E*. sp.1 originally identified as *E. klossi*, while the two specimens of *E.* sp.2 were identified individually as *E. micrura* and *E. grandis* ^[91]^. However, the *CYTB* gene trees of *Euroscaptor* support our finding that these specimens instead belong to cryptic lineages that each form a well-supported monophyletic clade, a conclusion also evidenced by the ABGD and ASAP analyses (**Fig. S3B**). Both lineages are moreover geographically separated from other species in this genus, with *E.* sp.1 distributed between the Salween and Mekong rivers of southwestern Yunnan, and *E*. sp.2 only being known from one locality to the west of the Salween River in southwestern Yunnan (**Fig. S4B**).

To further validate these putative species, we conducted multispecies coalescent analyses using 11 nuclear genes, as implemented in Bayesian Phylogenetics and Phylogeography (BPP).

Due to insufficient taxon sampling, however, *Euroscaptor* sp.2 was not included in the BPP analysis. In line with the *CYTB* results, BPP employing various combinations of algorithms and parameters consistently supported the genetic distinctiveness of *Parascaptor* sp.1 and *P.* sp.2, as well as supporting *Euroscaptor* sp.1 as a well-defined species (**Table S3**).

Notably, our *CYTB* and species delimitation analyses also uncovered distinct subclades within several recognized *Euroscaptor* species (e.g., *E. orlovi*, *E. kuzentsovi*) and revealed that *E. parvidens* is paraphyletic. Accordingly, species diversity within this genus is almost certainly larger than currently described.

### 3.3 Measurement- and outline-based morphometric analyses support subtle but consistent differences between recognized and cryptic species

To complement the above molecular delimitation results, we examined the morphological distinctiveness of *Parascaptor* sp.1, *P*. sp.2, and *Euroscaptor* sp.1 using a morphometric analysis with 15 cranial measurements and a geometric morphometric analysis based on the outline shape of the ventral view of the skull. For *Parascaptor*, principal component analysis (PCA) based on both linear measurements and outline shapes failed to clearly separate *P. leucura* and the two putative species (**Fig. S5A, E**), indicative of a high degree of morphological similarity. For instance, in the measurement-based analysis, a plot of the first two principal component scores (PC1 vs. PC2, respectively) shows that *Parascaptor* sp.2 is generally plotted more negatively on the PC2 axis and more positively on the PC1 axis. Although this positioning allows for a marginal separation from *P. leucura* and *P.* sp.1, plots of the latter two species overlap in the positive side of PC2. PC2 accounts for 14.4% of the variation and is positively correlated with height of coronoid process (HCP) (loading >0.89), and negatively correlated with lower molar row length (LLM) (loading <-0.52) (**Table S4A**). This indicates that *P. leucura* and *P*. sp.1 are characterized by longer postpalatal length, wider cranial breadth, but a shorter lower molar row, compared to *P.* sp.2. On the other hand, canonical variate analysis (CVA) successfully differentiated all three species with 100% accuracy, with all three species being well separated in the plots of the first two canonical variates (**Fig. S5C, G**).

In the case of *Euroscaptor*, measurement-based PCA and CVA both showed that *Euroscaptor* sp.1 plots closely with *E. klossi*, while the outline-based PCA revealed a close association of *E.* sp.1, *E. klossi*, and *E. micrura* (cf. **Figs. S5B, D** with **Fig. S5F**). In the PCA plot of the measurement-based analysis, *E.* sp.1 is plotted in the negative region of PC1 and the positive region of PC2, suggesting an overall smaller skull (PC1 is positively correlated with all measurements), a larger distance between the upper molars, and a broader foramen magnum breadth (**Table S4B**). This distinctiveness is supported by the outline-based CVA analysis which clearly differentiated *Euroscaptor* sp.1 from *E. klossi*, *E. micrura* and the two other closely related species included in the analysis. In summary, while PCA support overall morphological conservatism among *Euroscaptor* and *Parascaptor* species, the CVA analysis reveals subtle and consistent differences that discriminate species and putative species, supporting the recognition of these putative species as distinct taxonomic entities.

### 3.4 CNN network reach human-expert level accuracy in species identification

As traditional cranial measurement methods may overlook morphological differences that distinguish genetically distinct talpid lineages, our final aim was to evaluate the potential of deep learning techniques for this purpose. Our final data set included 2998 cranial photos from 819 specimens (generally four different views – ventral skull, dorsal skull, lateral skull, and lateral mandible – per specimen) representing 51 species/putative species across 18 genera (**Table S5**). Prior to building a convolutional neural network (CNN) for species identification, we first compared the identification accuracy of 19 different network models at the genus level using a 224x224 pixel image data set to identify the most suitable CNN architecture for this task. The networks were initially trained on a subset (∼80%) of photos (data set Trn0-224; **Table S6**) followed by testing on the remaining ∼20% (data set Tst0-224), resulting in genus identification accuracies ranging from 43.7-92.9% (**Table S7**). We then applied various photo manipulation techniques (**Fig. S12**) to create an augmented 21,429 photo data set (Trn1-224) which was shown to improve network identification accuracy by 3.2%-39.5%. As the EfficientNet series exhibited a moderate computational efficiency yet was found to consistently achieve the highest identification accuracy on both the non-augmented and augmented data sets, we thus further explored the impact of image resolution on genus identification accuracy when using this model series. Identification accuracy was found to progressively increase at higher image resolutions, with all EfficientNet models achieving accuracies >96.5% at 380x380 pixels (**Table S8**). Accordingly, we selected the EfficientNet-B3 model with a 380x380 pixel resolution for subsequent analyses based on its superior accuracy (98.0%) and workable computational efficiency.

We next tested the effects of integrating different cranial views and a hierarchical identification strategy for species identification accuracy. When we first attempted to match single specimen images – either the lateral mandible view or the lateral, ventral, or dorsal views of the skull – to one of the 51 species (one-step strategy), 27 species (52.9%) were always correctly identified (**Fig. S6**), with overall species identification accuracy being 88.0% (**Table 1**). Reconciling the results of up to four images per specimen increased the number of species (37 of 51 or 72.5%) always corrected identified (**Fig. S7**), though the average species identification accuracy was largely unchanged (88.5%).

**Table 1.**
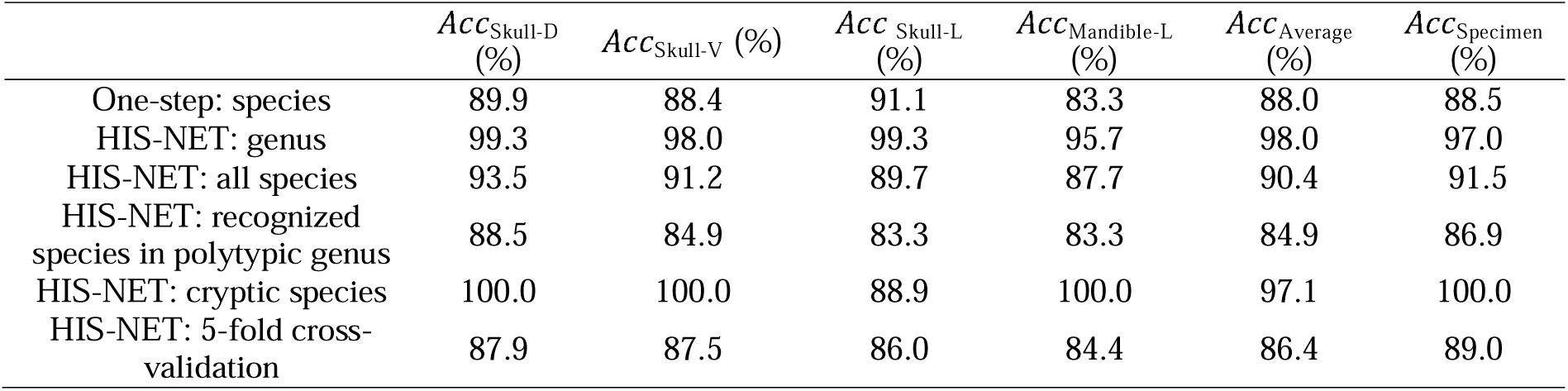
A summary of species identification (Acc) percentages (%) in different analyses based on the images of lateral (_Skull-L_), ventral (_Skull-V_), and dorsal (_Skull-D_) views of the skull and the lateral mandible view (_Mandible-L_).

When implementing a hierarchical identification strategy (**Fig. 2**), the average single image-based identification accuracy at the genus level was 98.0% with 12 of 18 (66.7%) genera always correctly identified (**Fig. S8**), while the multi-photo specimen-based analysis always correctly identified 14 genera (77.8%) with a mean accuracy of 97.0% (**Fig. S9**). At the species level, the mean image-based and specimen-based identification accuracies were slightly lower at 90.4% and 91.5%, respectively (**Table 1**). We further evaluated the robustness of our network using a five-fold cross-validation strategy (**Table S9**). This resulted in an image-based accuracy of 86.4% and a specimen-based accuracy of 89.0%. In conclusion, the hierarchical approach enhanced the accuracy by ∼2-3% and reached close to 90% specimen-based accuracy, demonstrating that our CNN network attained a human-expert level accuracy. We named our network HIS-NET (Hierarchical Identification of Species Network).

**Figure 2:**
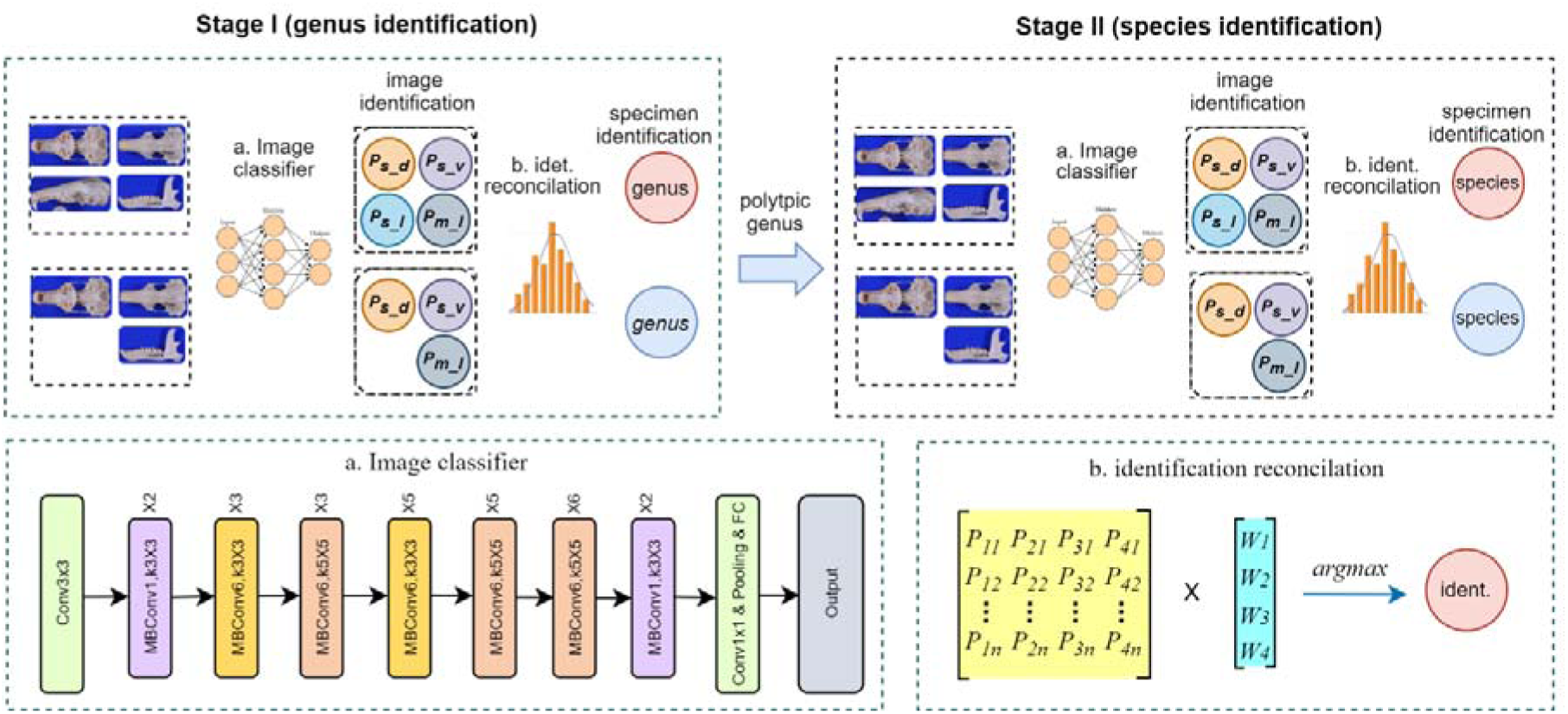
The architecture of HIS-NET (Hierarchical Identification of Species NETwork). Digital photographs of multiple skull views (skull dorsal view, s_d; skull ventral view, s_v; skull lateral view, s_l; mandible lateral view, m_l) were first processed for each specimen by CNN-based classifiers (a) to obtain a probability matrix (P) per image. Image reconciliation (b) was then performed to sequentially predict the genus (Stage I) and species (Stage II) of the specimens. The lower left panel illustrates the image classifier’s structure, comprising multiple convolutional, pooling, and fully connected layers (see text for details). The lower right panel depicts the reconciliation process, where the probability matrix (P) for each specimen’s images is multiplied with a view-specific weight vector (W) to produce the final identification through an argmax operation.

It is worth stressing that the identification of small mammal specimens often presents additional challenges due to the frequent occurrence of missing or broken skull and/or mandible elements. These conditions not only complicate identification efforts but often necessitate the exclusion of such specimens from both morphometric and geometric morphometric analyses. We thus tested the accuracy of species identification across various combinations of image availability and specimen integrity (**Table S9**). Our findings indicate that when four images per specimen were available, or when three images included at least two complete views, the HIS-NET model consistently achieved species identification accuracies ≥ 88%. Conversely, when two out of three views of the skull were broken, or when the total number of images was limited to one or two, the identification accuracy dropped below 75% (**Fig. S10**). These results underscore the need for careful consideration of specimen condition and the development of standardized imaging protocols in the implementation of such technologies.

### 3.5 HIS-NET accurately identifies cryptic species and reveals novel diagnostic characters

Our final goal was to examine whether the genetically identified cryptic species could be distinguished correctly from their con-generic partner. The three cryptic species included in this analysis (*E.* sp.1, *P.* sp.1, *P.* sp.2) were correctly identified with 100% accuracy (**Fig. 3**), which is higher than the mean identification accuracy of recognized species within the seven studied polytypic talpid genera (86.9%). This finding underscores the ability of HIS-NET to efficiently distinguish cryptic species based solely on cranial photos.

**Figure 3.**
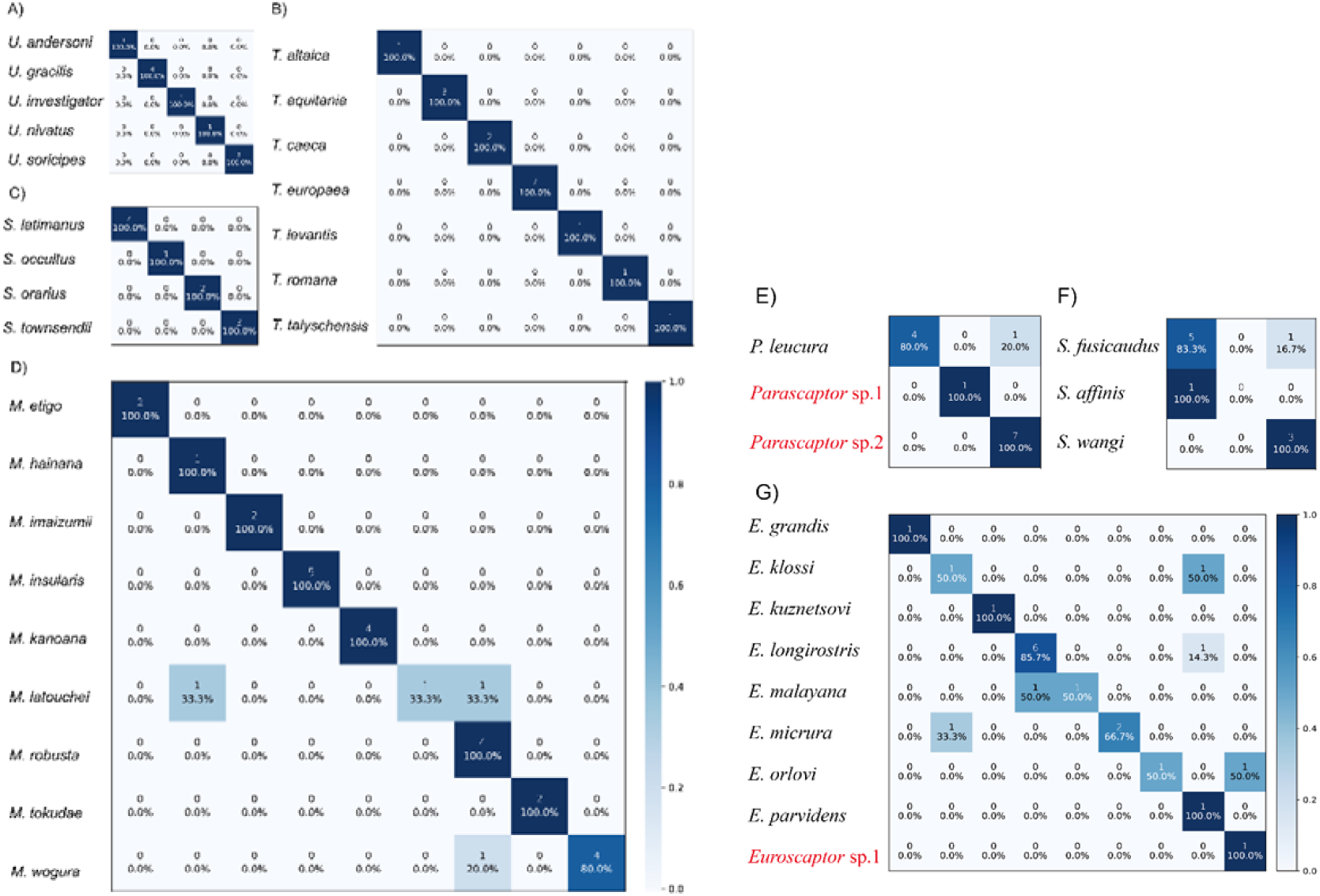
Confusion matrix heatmap showing HIS-NET identification accuracy for recognized and cryptic species in all polytypic talpid genera. The panels illustrate the accuracy for (A) *Uropsilus*, (B) *Talpa*, (C) *Scapanus*, (D) *Mogera*, (E) *Parascaptor*, (F) *Scaptonyx*, and (G) *Euroscaptor*. Numbers at the top of each cell denote the number of specimens assigned to each species.

To better understand the model’s decision-making process, we examined Class Activation Mapping (CAM) generated heatmaps to identify morphological features that HIS-NET deemed most informative for species discrimination. Across all taxonomic groups and regardless of whether the focus was on the skull or mandible, HIS-NET consistently prioritized dental regions, particularly the premolars and molars (**Fig. 4**). For instance, the upper premolars P4 exhibited notable variations across the three species/putative species of *Parascaptor*: in *P. leucura*, P4 is characterized by a well-developed paracone with a crest extending posteriorly that connects with a low metastyle, forming a near-triangular shape on buccal view. The P4 paracone of *P.* sp.1 extends from a crest that descends vertically without directly connecting to the metastyle cusp. The P4 morphology of *P.* sp.2 is reminiscent of its P2, with a crest extending posteriorly from the paracone but not directly connecting to the more posteriorly positioned metastyle, resulting in a trapeziform shape of P4 when viewed buccally. However, the developmental progression of P4 follows a different order: *P.* sp.2 > *P. leucura* > *P.* sp.1, consistent with the topology of the *CYTB* gene tree.

**Figure 4.**
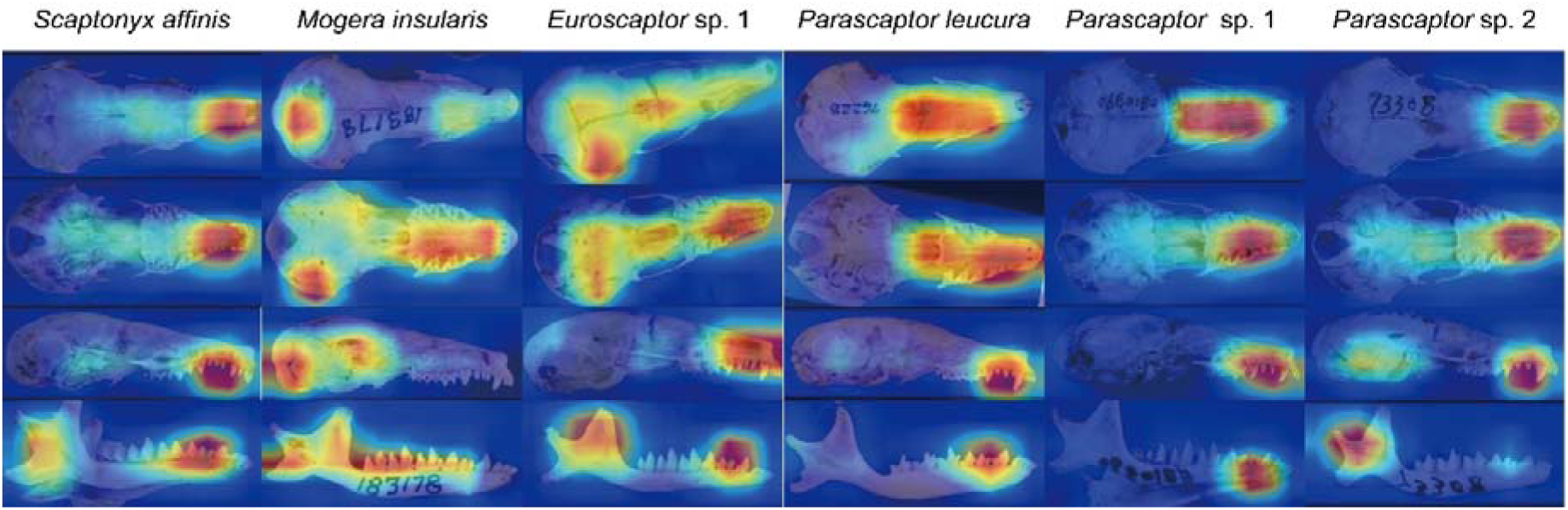
Example of class activation mapping (CAM) heatmaps obtained from representative talpid species and putative species. For each species, the images in each column from top to bottom represent the dorsal, lateral and ventral views of the skull and the lateral view of the mandible, respectively. The relative importance of each skull region that the model uses for species identification is denoted by color, with red and blue areas being of high and low importance, respectively. The images show that the dental regions, as well as palatine, pterygoid, and tympanic regions are important contributors for the model’s decision-making process.

For the skull, the model also focused on the maxillary region in both ventral and lateral views, as well as the nasal and frontal areas in the dorsal view (**Fig. 4**). Additionally, HIS-NET frequently highlighted the palatine, pterygoid, and tympanic regions, with occasional emphasis on the parietal bone in some specimens. Most of these areas have long been known to be critical for species identification in small mammals. In the case of the mandible, the model consistently emphasized the ascending ramus, with particular attention to the area surrounding the condylar process. The observation that HIS-NET consistently focused on known informative anatomical regions emphasize the model’s ability to detect specific features that are crucial for species identification. However, our examination of CAM-highlighted areas in *Euroscaptor* revealed additional intriguing features beyond differences in upper and lower dental morphology. Notably, CAM highlighted the basipharyngeal region as crucial for species identification of *Euroscaptor* sp.1. The pterygoid hamuli, which form the walls of the basipharyngeal canal in this species, extend laterally on the anterior half. This feature is distinctive among all examined *Euroscaptor* species and has not previously been utilized as a diagnostic character in talpid taxonomy. These findings emphasize the power of CAM in assisting the identification of novel, effective diagnostic features that traditional morphometric and geometric morphometric approaches may overlook.

## 4. Discussion

### 4.1 New hypothesis of Talpidae evolution

Our phylogenetic analyses using UCEs corroborate and are largely congruent with previously published topologies derived from Tree of Life genes ^[8, 37]^. Briefly, all three studies fail to support the monophyly of the two strictly fossorial tribes (Scalopini and Talpini) thereby supporting the contention first suggested by Shinohara et al. ^[92]^ that this derived lifestyle evolved twice independently in this family. Desmanini was also recovered as the sister group to a clade containing shrew moles, star-nosed moles, and Eurasian fossorial talpids congruent with the concatenation analysis of He et al. ^[37]^. Similarly, the UCE concatenation analysis placed Chinese shrew moles (*Scaptonyx*) sister to North American shrew moles (*Neurotrichus*) in line with both Tree of Life studies ^[8, 37]^. The only major discordance among these studies pertains to the placement of *Condylura*, which has been notoriously difficult to resolve ^[92]^. Importantly, the current analyses reveal a novel and well-supported sister relationship between the star-nosed mole (*C. cristata*) and shrew moles. This finding conflicts with earlier studies that placed *Condylura* sister to Desmanini ^[8]^ or Talpini ^[37]^ with moderate to low support. Accordingly, our robust phylogenetic hypotheses provide a valuable framework for further investigation into the systematics of talpids, encompassing both extant and extinct lineages. Importantly, the results support the presence of multiple undescribed lineages within widespread recognized species or species complexes, highlighting the rich species diversity in the sky islands of Southwest China.

### 4.2 Unraveling the underestimated species diversity of talpid mole

Our comprehensive study, combining ultraconserved element (UCE) phylogenomics, molecular species delimitation, traditional morphometrics, and cutting-edge CNN-based analyses, reveals a previously underappreciated diversity within the genera *Euroscaptor* and *Parascaptor*. Each cryptic lineage identified in the UCE phylogeny emerged as a monophyletic clade and is supported by measurable and diagnosable morphological characters despite the overall morphological conservatism within each genus. These cryptic species are moreover confined to restricted mountain ranges (**Fig. S4**) and conform to IBD/IBE diversification principles. Collectively, these lines of evidence unambiguously support the recognition of these cryptic lineages as full species, even under the stringent criteria of the dmPSC.

Our findings indicate that *Parascaptor* comprises at least three distinct species: *P. leucura*, distributed across Nepal, northeastern India, and parts of western Myanmar; *P.* sp.2, inhabiting central and western Yunnan, southwestern Sichuan, and eastern and central Myanmar; and *P.* sp.1, found in central-eastern and northeastern Yunnan and adjacent southern Sichuan. Although genetic data for *P. leucura* were not included in this study, unpublished data (pers. comm. Shi-Ichiro Kawada) indicate that *Parascaptor* sp.1 is sister to a clade comprising *P. leucura* and *Parascaptor* sp.2. This phylogenetic relationship aligns with our morphological analyses, which reveal that *Parascaptor* sp. 1 exhibits the most primitive dental (current study) and mandibular ^[34]^ morphology of the three species.

In *Euroscaptor*, our analyses support the recognition of *Euroscaptor* sp.1 from Menglian. Originally identified as *E. klossi* following their collection in 1964, these specimens instead form a lineage distinct from other nearby distributed species (*E. klossi, E. kuznetsovi*, and *E. orlovi*) (**Fig. S4**). The status of *Euroscaptor* sp.2, represented by two samples (one with a broken skull and skin and the other with only skin) from Yingjiang, remains less certain. While mitochondrial data clearly differentiate it from *E. grandis*, we cannot rule out a potential affinity to *E. micrura* without further phylogenetic analysis including this species.

The phylogeographic patterns observed in our newly identified talpid mole species provide compelling evidence for the role of sky islands in driving allopatric diversification across Southwestern China and adjacent mountain regions. Our findings reveal a clear IBD and IBE diversification pattern (**Fig. S4**), with species distributions confined to middle-to-high elevation habitats within distinct mountain ranges that are separated by large valleys and drainage systems. This pattern of distribution is consistent with observations in other animal groups inhabiting this region ^[17–21]^, underscoring the generality of these biogeographic processes.

Importantly, our study unveils a hierarchical clustering pattern that adds a layer of complexity to the observed diversification. For instance, within *Parascaptor* sp.2, we identified distinct lineages from central Myanmar and western Yunnan (west of the Salween River), with each forming a separate clade/lineage (**Fig. S3**). This genetic structuring suggests ongoing diversification processes. Similarly, our analysis of the *CYTB* gene tree revealed that *Euroscaptor parvidens* comprises two non-monophyletic clades (**Fig. S3**), indicating an underestimation of species diversity within this taxon in the mountains of central and southern Vietnam. The finding of potentially distinct lineages in *Euroscaptor orlovi* and *E. kuzentsovi* additionally emphasize the need for further investigation with increased sampling efforts to fully elucidate the extent of cryptic diversity in southeast Asia.

### 4.3 Advantage of CNN in species identification

While CNNs have revolutionized species identification in various taxa like plants and insects ^[46–48]^, their application in mammalian taxonomy, particularly for small mammals, has remained largely untapped. This is largely due to a lack of comprehensive skull and dental image databases for any group of mammals due to the high cost and time investment required for data collection. Species sample sizes of available museum specimens are also uneven, which is especially true for rare and recently recognized species. Finally, training CNNs to recognize minute interspecific differences in skull and dental features is a formidable task. To overcome these challenges, we first constructed global-scale image database encompassing nearly all genera and a majority of species within the talpid family. We also adopted an augmentation strategy which helped to balance the different sample sizes between genera and species. The high levels of species identification accuracy demonstrate the potential of CNNs for distinguishing (and discovering) nuanced morphological differences and highlights the potential applicability of deep learning networks using cranial images to other groups of mammals.

Our HIS-NET, employing a hierarchical approach with sequential genus and species-level classifications, achieved an impressive overall accuracy exceeding 98% at the genus level and 91% at the species level, which represents an improvement of 2-3% over the one-step identification strategy. Direct species identification within our dataset would require processing an average of 6.0 bits of Shannon information entropy per identification, whereas our hierarchical approach reduced this to 4.2 bits for genus-level and 2.8 bits for species-level classifications. This suggests that by breaking down the identification task, the network requires less information at each stage, making species decision more efficient. Moreover, hierarchical classification aligns with traditional taxonomic practice, where genus-level distinctions are often more conspicuous than subtle interspecific differences. By first establishing a strong genus-level classification, HIS-NET likely benefits from a reduced search space and can focus on finer-scale morphological distinctions for species-level identification. Future research could explore adding subfamily and tribe hierarchical layers to potentially enhance computational efficiency and identification accuracy even further.

The robustness of HIS-NET in identifying species from specimens with damaged or missing features that are typically not included in traditional and geometric morphometric analyses is particularly noteworthy. This capability, enhanced by our data augmentation strategy simulating cranial damage (e.g., drop out and coarse drop out), addresses a common challenge in taxonomic studies of small mammals where specimens are often incomplete. However, specimens with fewer images combined with a higher percentage of broken skulls/mandibles were more prone to misidentification (**Fig. S10**).

One of the most exciting aspects of our CNN approach is its ability to reveal previously overlooked morphological characters. For instance, the network found that the structure of the basipharyngeal canal in *Euroscaptor* species is highly effective in distinguishing between species of this genus (Fig S11). Analysis of this structure has been employed for other groups of mammals ^[93]^ though has previously been overlooked in Talpidae taxonomy. This discovery thus underscores the potential of CNNs in guiding taxonomists towards a novel and valuable repertoire of distinguishable morphological characters for species delimitation, which in turn could accelerate our ability to discern cryptic species more efficiently.

Our results also illuminate several avenues for future research and methodological improvements. First, because we show that incorporating different views of the skull improves identification accuracy, expanding the image database to include more views, such as the occlusal surfaces of mandibles, and/or taking multiple photos from different angles of the same view could further improve accuracy ^[47, 94]^. Indeed, the occlusal view includes dental morphology that is crucial for identification of extant and fossil species. Second, while our CNN achieved expert-level accuracy, the CAM revealed that the network tends to focus on specific regions of each image. This selective attention, while effective, suggests that our model has not yet fully captured the complexity of morphological variation present in the skulls and teeth of talpid moles. Thus, incorporating advanced machine learning techniques such as attention mechanisms ^[95]^ or object detection algorithms ^[96]^, coupled with specific guidance such as labeling individual teeth and auditory bullae, could further enhance the model’s accuracy. Third, an intriguing finding of our study was the unexpectedly high accuracy rates achieved from the dorsal skull view, which were comparable to those obtained from ventral and lateral views (**Table 1**). This observation challenges the notion that dorsal skull features are less informative for species identification in vertebrates. It further suggests that there may be previously unrecognized or underappreciated morphological characters in the dorsal skull region that are taxonomically significant.

In conclusion, our CNN-based approach represents a technological advancement in mammalian taxonomy, offering a powerful tool for bridging the gap between molecular species delimitation and traditional morphological examination. It also highlights the potential for further refinement and expansion of these methods. The integration of deep learning with traditional taxonomic methods promises to revolutionize our ability to discern and discover species, particularly in groups with subtle morphological differences.

## Conflict of interest

The authors declare that they have no conflict of interest.

## Acknowledgements

We thank the curators and staff at the Smithsonian Institution, National Museum of Natural History (KM Helgen, D Lunde), Field Museum (L Heaney), Burke Museum (B Sharon) and New Mexico Museum of Natural History (B Oh), Kunming Institute of Zoology (S Li) and Sichuan Academy of Forestry (R Liao) for approving our proposal for the use of tissue samples, or conducting destructive sampling on museum skin specimens. We also thank museum curators and staff at the Guangdong Institute of Zoology (JH Zheng), National Museum of Natural Science at Taichung (YJ Chen), American Museum of Natural History (NB Simmons, E Westwig), National Museum of Natural History (KM Helgen, D Lunde), Field Museum (L Heaney), Museum of Comparative Zoology (HE Hoekstra and J Chupasko), National Museum of Nature and Science of Japan (S-I Kawada), Hokkaido University Natural History Museum (T Tsubota), Institute of Ecology and Biological Resources of Vietnam (TS Nguyen), the State Museum of Natural History Stuttgart (E Amson, C Leidenroth, S Merker) and the University of Montpellier (PH Fabre) for allowing us to examine specimens and take photos of specimens under their curation. We thank J Decher, C Montermann for kindly providing us photos of *Galemys*, and T Martin, C Steinweg for kindly providing us photos of *Desmana*. We also thank R Haslauer for extensive discussion of the taxonomy of Talpidae. We are grateful to Umi Matsushita for painting the talpid species used in the figures. This work was supported by the National Natural Science Foundation of China (32170452), GuangDong Basic and Applied Basic Research Foundation (2022B1515020033), Key Program of the National Natural Science Foundation of China Regional Innovation and Development Joint Fund (U23A20161) and the Open Project of Ministry of Education Key Laboratory for Ecology of Tropical Islands, Hainan Normal University (HNSF-OP-2024-3) to KH, the National Science Foundation (NSF DEB-1457735 to MSS), and by the University of Manitoba Research Grants Program (41342), National Sciences and Engineering Research Council of Canada Discovery (RGPIN/238838–2011; RGPIN/6562–2016) and Discovery Accelerator Supplement (RGPIN/412336–2011) grants to KLC.

## Author contributions

ZC and KLC designed and supervised the project. KH, QM, and AS photographed the specimens. ALL and KHW performed deep learning analyses. KH, ZZC, WYS, WHY, SYL, SL, and FYT conducted fieldwork and collected samples. KH, XYW, JYL, and MS analyzed genetic data. KH and SWH performed molecular laboratory experiments. KH, SNX, ZLZ, ZQY, and HR conducted morphometric analyses. ZLZ, SYL, QQL, and XYZ carried out data annotation for deep learning. HBY performed GIS analyses.

## Supplementary materials

### Supplementary figures

**Figure S1.** Phylogenetic tree of Eulipotyphla (Talpidae, Soricidae, Erinaceidae and Solenodontidae) based on a concatenated alignment of ultraconserved elements. Branch lengths represent substitutions per site. Unless specified, all relationships are highly supported (bootstrap value =1.0).

**Figure S2.** Phylogenetic tree of Eulipotyphla (Talpidae, Soricidae, Erinaceidae and Solenodontidae) based on a coalescent analysis of ultraconserved elements. Branch lengths represent coalescent units. Unless specified, all relationships are highly supported (bootstrap value =1.0). Dashes indicate weakly supported relationships (bootstrap value < 0.5), while the hashtag indicates a relationship different from that estimated in the concatenation analysis.

**Figure S3.** A heatmap showing the Kimura 2-parameter distance (K2P) of the *CYTB* gene among species and samples in (A) *Parascaptor* and (B) *Euroscaptor*. *CYTB* gene trees for each genus are shown to the left and top of each heatmap. Vertical black bars represent species delimitation results from the Automatic Barcode Gap Discovery (ABGD) and Assemble Species by Automatic Partitioning (ASAP) analyses. The vertical blue bars to the right indicate taxonomic affiliations. Displayed ultrametric trees are maximum likelihood gene trees estimated using complete *CYTB* sequences, as implemented in RAxML (see text for details). Results demonstrate that specimens in *Parascaptor* sp.1 and *P.* sp.2 each form reciprocally monophyletic clades, while specimens in *Euroscaptor* sp.1 and *E*. sp.2 also form distinct clades. Note that *E. parvidens* is shown to comprise two non-monophyletic clades in all three analyses.

**Figure S4.** Distribution map of recognized and putative species in (A) *Parascaptor* and (B) *Euroscaptor* (B).

**Figure S5.** Results of principle component analysis (PCA) and canonical variate analysis (CVA) based on measurement-based morphometric and outline-based geometric morphometric analyses. A-D: PCA plots showing scores on PC1 and PC2 derived from 15 log_10_-transformed craniomandibular variables for (A) *Parascaptor* and (B) *Euroscaptor*, and corresponding CVA plots displaying scores on CV1 and CV2 for the same data in (C) *Parascaptor* and (D) *Euroscaptor*. E-H: PCA plots illustrating scores on PC1 and PC2 based on skull outline shapes for (E) *Parascaptor* and (F) *Euroscaptor*, and corresponding CVA plots showing scores on CV1 and CV2 for the same data in (G) *Parascaptor* and (H) *Euroscaptor*.

**Figure S6.** Confusion matrix heatmap showing the image-based identification accuracy for recognized and cryptic species across all species and putative species using a one-step strategy. Each photo was directly identified to one of the 51 species. The Y-axis (true_label) represents the actual species classification of each species, while the X-axis (predicted_label) represents the species classification result predicted by the model. Numbers at the top of each cell denote the number of specimens assigned to each species, while the percentage of specimens assigned to that species is listed at the bottom of the cell. The overall accuracy of the model, based on the correct identification of individual photos, was 88.0% (see Table 1).

**Figure S7**. Confusion matrix heatmap showing the individual-based identification accuracy for recognized and cryptic species across all species and putative species using a one-step strategy. Each specimen was directly identified to one of the 51 species. The Y-axis (true_label) represents the actual species classification, while the X-axis (predicted_label) represents the species classification result predicted by the model. Numbers at the top of each cell denote the number of specimens assigned to each species, while the percentage of specimens assigned to that species is listed at the bottom of the cell. The overall accuracy of the model, based on the correct identification of individual specimen, was 88.5% (see Table 1).

**Figure S8.** Confusion matrix heat map showing the image-based identification accuracy for all genera using HIS-NET. The Y-axis (true_label) represents the actual species classification, while the Y-axis (predicted_label) represents the species classification result predicted by the model. Numbers at the top of each cell denote the number of specimens assigned to each species, while the percentage of specimens assigned to that species is listed at the bottom of the cell. The overall accuracy of the model, based on the correct identification of individual images, was 98.0% (see Table 1).

**Figure S9.** Confusion matrix heatmap showing the specimen-based identification accuracy for all genera using HIS-NET. The Y-axis (true_label) represents the actual species classification, while the X-axis (predicted_label) represents the species classification result predicted by the model. Numbers at the top of each cell denote the number of specimens assigned to each species, while the percentage of specimens assigned to that species is listed at the bottom of the cell. The overall accuracy of the model, based on the correct identification of individual specimens, was 97.0% (see Table 1).

**Figure S10.** Summary of species identification accuracy obtained following five iterations of cross-validation with HIS-NET. Each bar corresponds to a specific category, with values below each bar indicating the number of available images per specimen, and those in parentheses representing the number of images for each specimen showing a broken skull or mandible element. The height of each bar reflects the total number of specimens per category, with the green and orange segments corresponding to the number of correctly and incorrectly identified specimens, respectively. Numbers above each bar indicate the total number of specimens examined per category with the overall percent species identification accuracy for each category given in parentheses.

**Figure. S11.** The lateral skull view of (A-C) three *Parascaptor* species/putative species and (D-F) the ventral skull view of three *Euroscaptor* species/putative species.

**Figure S12**. Illustration of the various augmentation strategies used in this study. The original image (top left) was subjected to different augmentation techniques to introduce minor distortions and mitigate overfitting during neural network training. The augmentation methods included noise injection (SaltAndPepper, AdditiveGaussianNoise, ImpulseNoise), blurring (GaussianBlur, AverageBlur, MotionBlur), rotation (Rot90, Affine (rotate 45 degrees)), mirroring (Flipud (vertical flip), Fliplr (horizontal flip)), masking (CoarseDropout, Dropout), and contrast adjustment (SigmoidContrast, GammaContrast and AveragePooling).

### Supplementary tables

**Table S1.** Taxonomy of Talpidae used in this study.

**Table S2.** Information regarding specimens used for UCE sequencing, Tree of Life (TOL) gene sequencing, and *CYTB* gene sequencing, as well as accession numbers of sequences downloaded from GenBank for species delimitation analyses.

**Table S3**: Summarized results of Bayesian Phylogenetics and Phylogeography (BPP) analyses for the genera (A) *Parascaptor* and (B) *Euroscaptor* using 11 nuclear genes.

**Table S4**. Results of PCA conducted based on fifteen craniomandibular variables for (A) Parascaptor and (B) Euroscaptor. Factor loadings, eigenvalues and percentage of variance explained for each principal component (PC) are shown.

**Table S5**. Summary of the number of specimens and photos used in the CNN-based analyses. A total of 51 species/putative species were included, following the taxonomy presented in Table S1. The images were divided into training and testing datasets with an approximate ratio of 4:1. Data augmentation was applied to the training dataset, with up to 30-fold augmentation for each image.

**Table S6.** Prepared datasets used in CNN image-based analyses. Images from the original data (Origin) were cropped (Origin-C) and padded to a square shape (Origin-CP). These images were then split into training and testing datasets with an approximate ratio of 4:1. The images were downsized for model selection (resolution 224 x 224) and resolution selection (224 x 224, 260 x 260, 300 x 300, and 380 x 380).

**Table S7.** Summary of parameters used in each model. The identification accuracy for each model using image data with and without augmentation is given (see text for details). Improvement is defined as the accuracy obtained using data with augmentation minus the accuracy without augmentation. The number of parameters (Params) and floating-point operations per second (FLOPs) for each model are also provided. Stochastic Gradient Descent (SGD) was used as Optimizer.

**Table S8**. Performance comparison (%) of the EfficientNet model series (B0, B2, B3 and B3) across various image input resolutions. The number of floating-point operations per second (FLOPs) required for each model are also provided.

**Table S9.** Summary of the results of the five-fold cross-validation of HIS-NET on a specimen identification basis. The table includes the number of specimens, the number of incomplete specimens (represented by fewer than 4 images), and the number of images used in the training and testing datasets, along with species identification accuracy (%). Additionally, it provides a summary of correctly (Corr.) and incorrectly (Incorr.) classified images for various scenarios based on the number of images and the extent of broken skulls/mandibles.

**Table S10**. Concatenation species tree estimated using (1) RAxML and (2) coalescent species tree estimated using ASTRAL-III. Branch length represents substitutions per site (concatenation) and coalescent units (coalescent).

## Appendix

Appendix I. Specimens used in image-based CNN analyses and measurement-based morphometric analyses

## Data availability

The data for genetic analyses was uploaded to Mendeley Data, V1, (doi: 10.17632/h32txw2xb9.1). The code of HIS-NET is available through https://github.com/Hua-jiu/HISNET

## Notes

### Competing Interest Statement

The authors have declared no competing interest.

