## Supplementary material for "Sky islands of Southwest China. II: Unraveling hidden species diversity of talpid moles using phylogenomics and skull-based deep learning": Fig. S

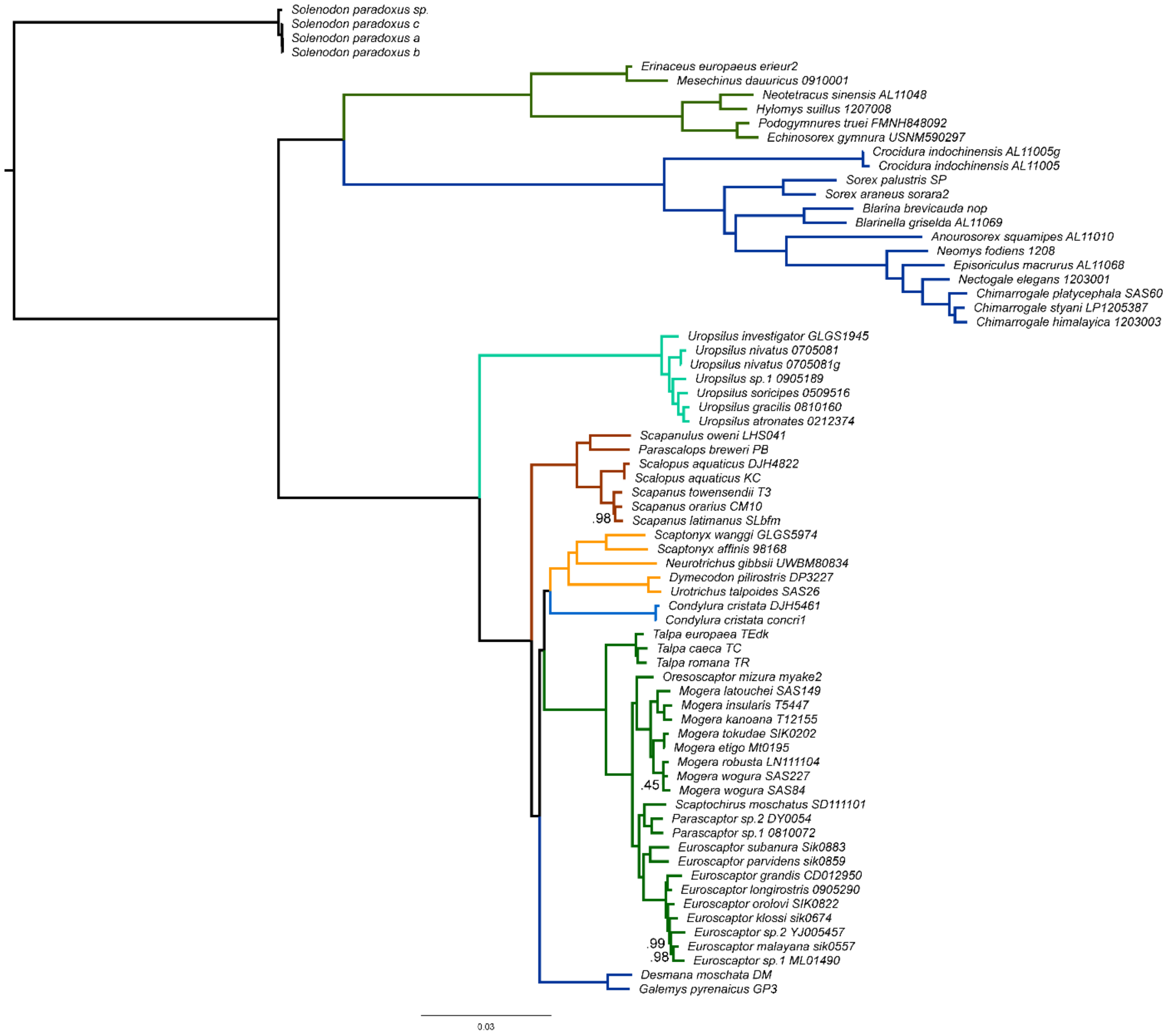


Figure S1. Phylogenetic tree of Eulipotyphla (Talpidae, Soricidae, Erinaceidae and Solenodontidae) based on a concatenated alignment of ultraconserved elements. Branch lengths represent substitutions per site. Unless specified, all relationships are highly supported (bootstrap value =1.0).

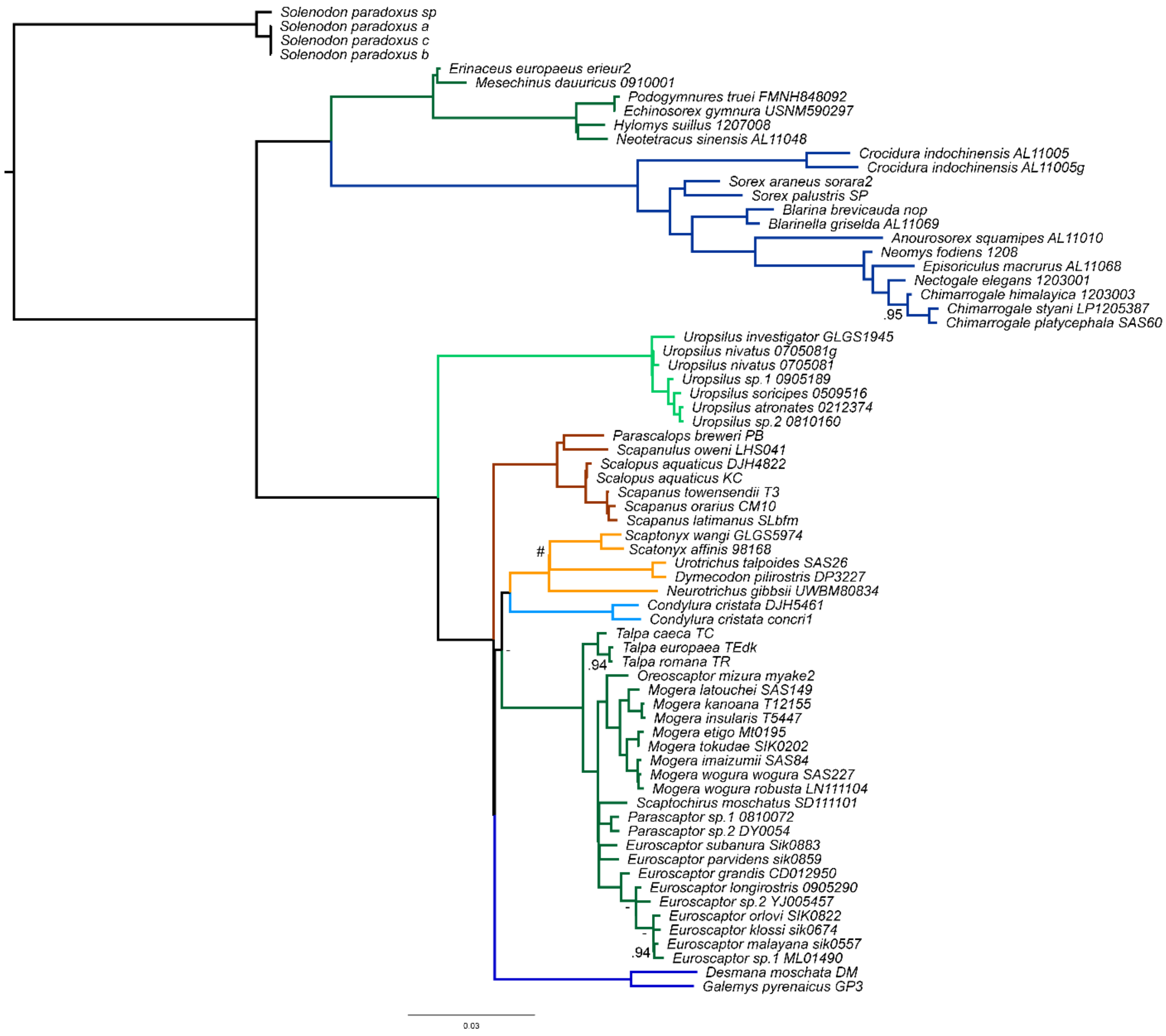


Figure S2. Phylogenetic tree of Eulipotyphla (Talpidae, Soricidae, Erinaceidae and Solenodontidae) based on a coalescent analysis of ultraconserved elements. Branch lengths represent coalescent units. Unless specified, all relationships are highly supported (bootstrap value =1.0). Dashes indicate weakly supported relationships (bootstrap value < 0.5), while the hashtag indicates a relationship different from that estimated in the concatenation analysis.

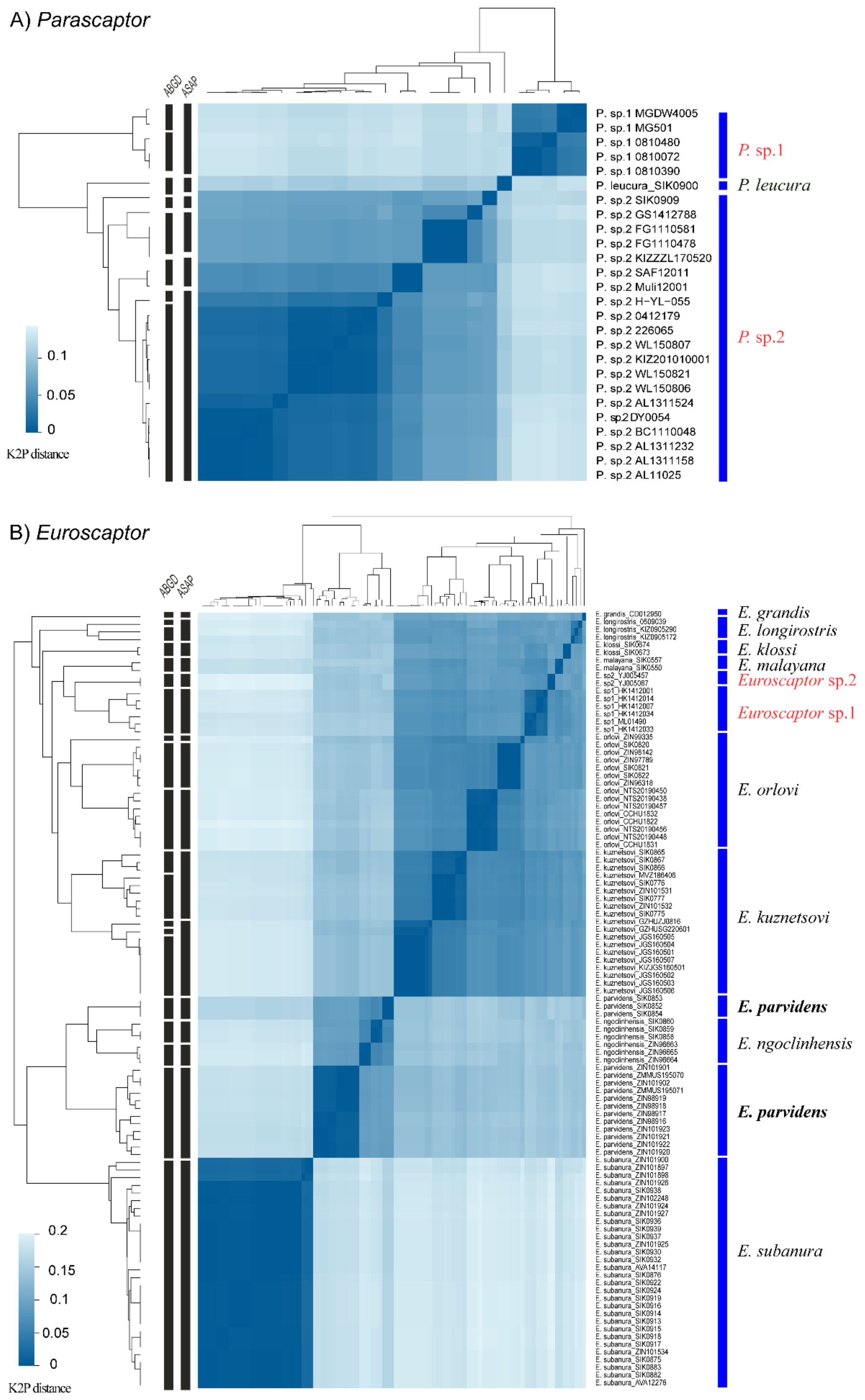

A) Parascaptor
ABGD
ASAP
P. sp.1 MGDW4005
P. sp.1 MG501
P. sp.1 0810480
P. sp.1 0810072
P. sp.1 0810390
P. leucura_SIK0900
P. sp.2 SIK0909
P. sp.2 GS1412788
P. sp.2 FG1110581
P. sp.2 FG1110478
P. sp.2 KIZZZL170520
P. sp.2 SAF12011
P. sp.2 Muli12001
P. sp.2 H−YL−055
P. sp.2 0412179
P. sp.2 226065
P. sp.2 WL150807
P. sp.2 KIZ201010001
P. sp.2 WL150821
P. sp.2 WL150806
P. sp.2 AL1311524
P. sp2 DY0054
P. sp.2 BC1110048
P. sp.2 AL1311232
P. sp.2 AL1311158
P. sp.2 AL11025
P. sp.1
P. leucura
P. sp.2
0.1
0.05
0
K2P distance
B) Euroscaptor
ABGD
ASAP
E. grandis
E. longirostris
E. klossi
E. malayana
Euroscaptor sp.2
Euroscaptor sp.1
E. orlovi
E. kuznetsovi
E. parvidens
E. ngoclinhensis
E. parvidens
E. subanura
0.2
0.15
0.1
0.05
0
K2P distance

Figure S3. A heatmap showing the Kimura 2-parameter distance (K2P) of the *CYTB* gene among species and samples in (A) *Parascaptor* and (B) *Euroscaptor*. *CYTB* gene trees for each genus are shown to the left and top of each heatmap. Vertical black bars represent species delimitation results from the Automatic Barcode Gap Discovery (ABGD) and Assemble Species by Automatic Partitioning (ASAP) analyses. The vertical blue bars to the right indicate taxonomic affiliations. Displayed ultrametric trees are maximum likelihood gene trees estimated using complete *CYTB* sequences, as implemented in RAxML (see text for details). Results demonstrate that specimens in *Parascaptor* sp.1 and *P.* sp.2 each form reciprocally monophyletic clades, while specimens in *Euroscaptor* sp.1 and *E*. sp.2 also form distinct clades. Note that *E. parvidens* is shown to comprise two non-monophyletic clades in all three analyses.

A) *Parascaptor* distribution

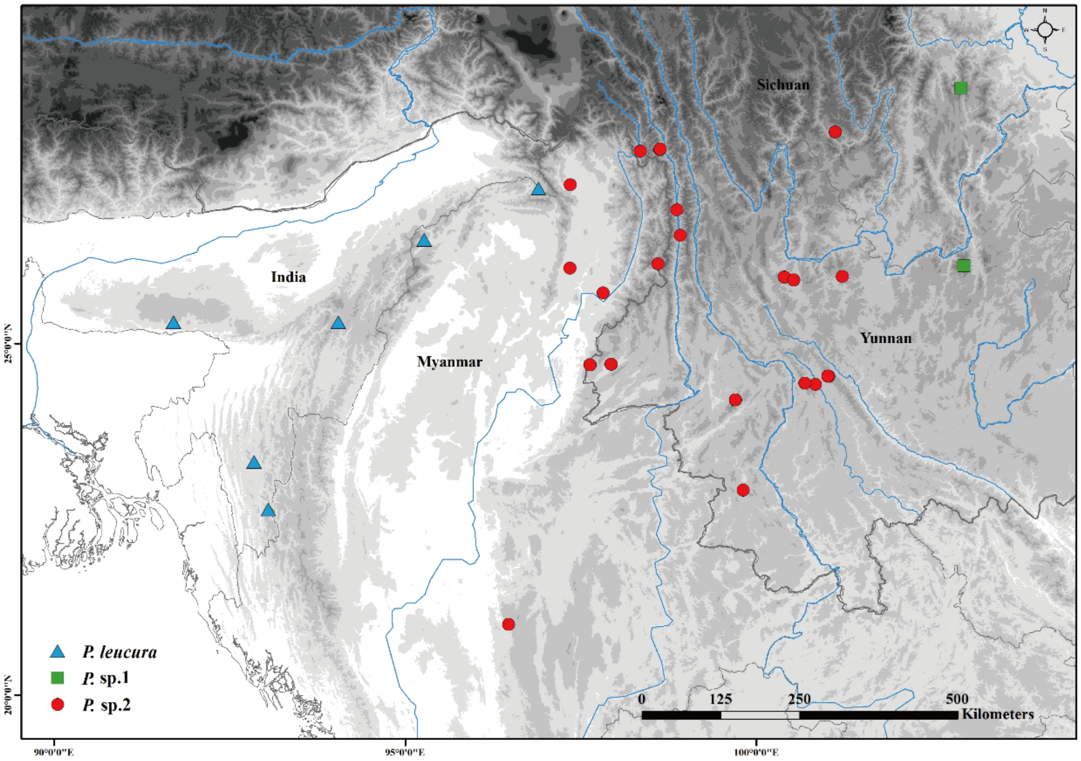


B) *Euroscpator* distribution

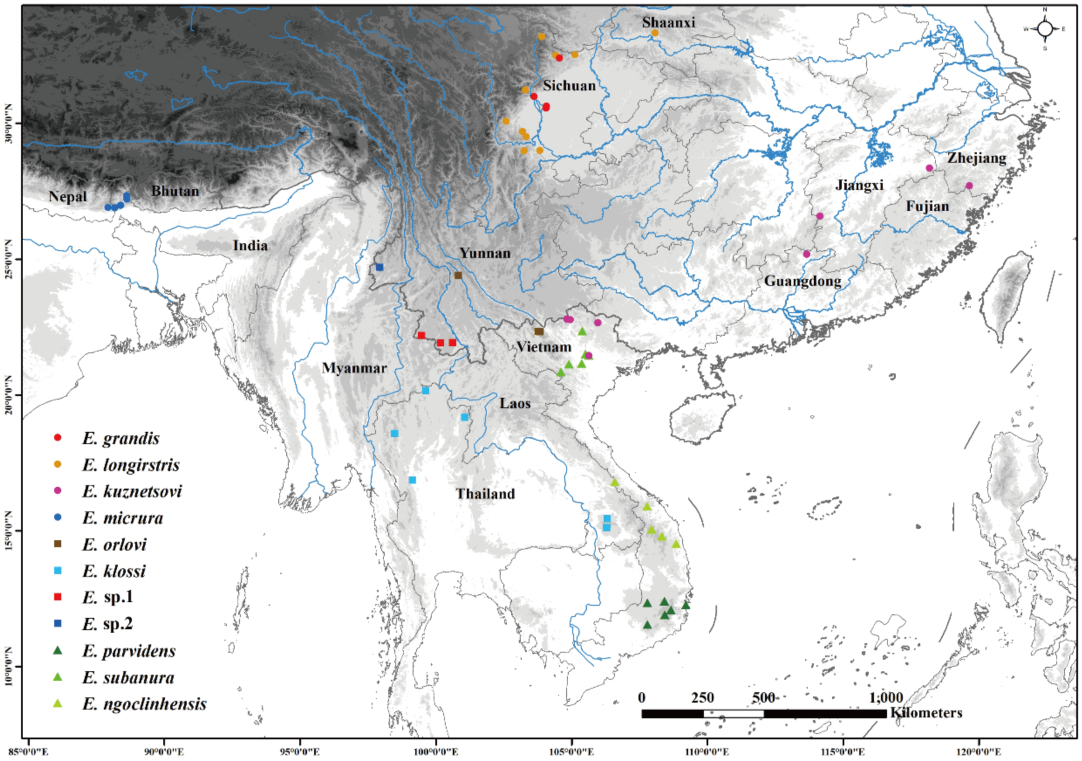


Figure S4. Distribution map of recognized and putative species in (A) *Parascaptor* and (B) *Euroscaptor*.

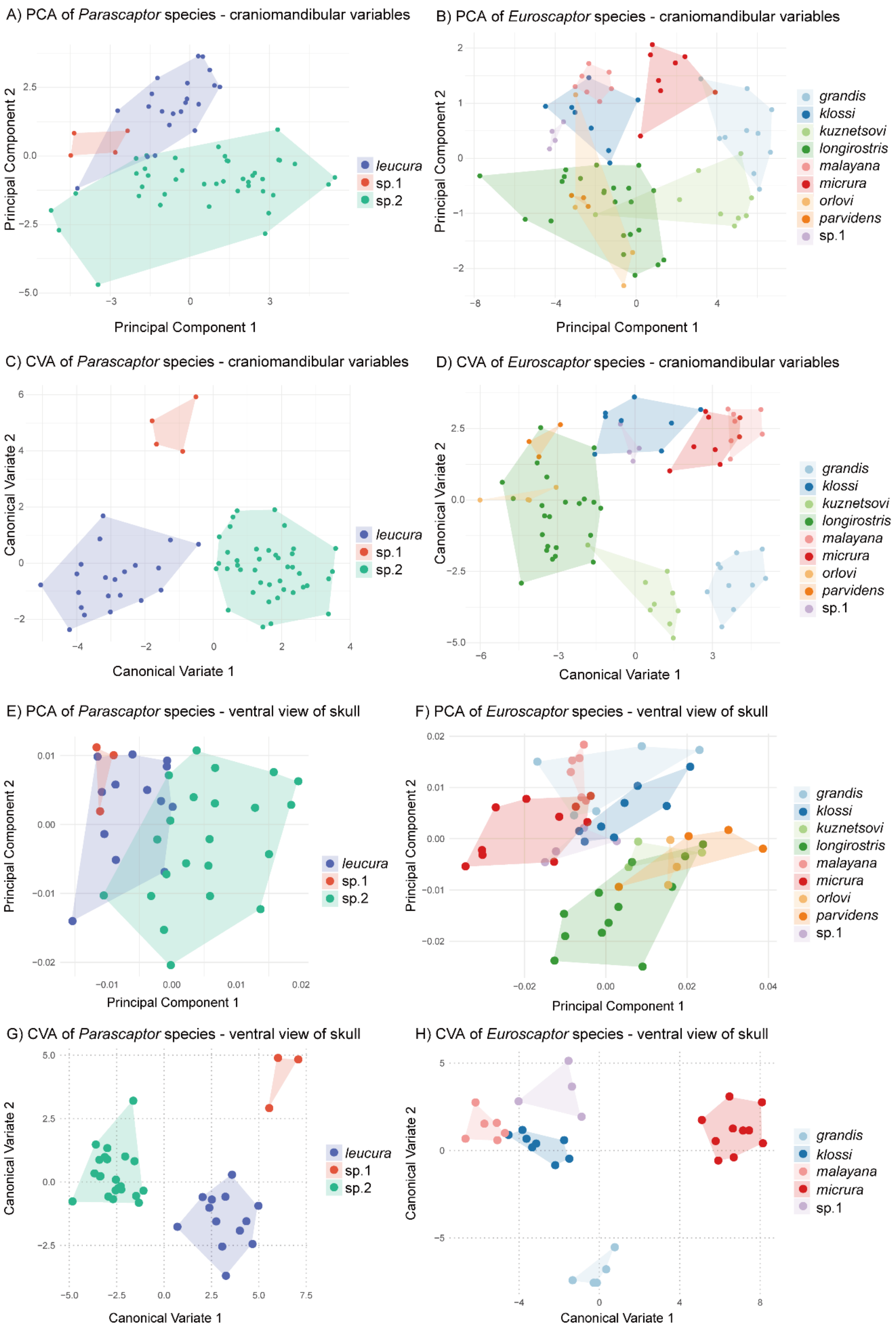

A) PCA of *Parascaptor* species - craniomandibular variables
Principal Component 1
Principal Component 2
*leucura*
sp.1
sp.2
B) PCA of *Euroscaptor* species - craniomandibular variables
Principal Component 1
Principal Component 2
*grandis*
*klossi*
*kuznetsovi*
*longirostris*
*malayana*
*micrura*
*orlovi*
*parvidens*
sp.1
C) CVA of *Parascaptor* species - craniomandibular variables
Canonical Variate 1
Canonical Variate 2
*leucura*
sp.1
sp.2
D) CVA of *Euroscaptor* species - craniomandibular variables
Canonical Variate 1
Canonical Variate 2
*grandis*
*klossi*
*kuznetsovi*
*longirostris*
*malayana*
*micrura*
*orlovi*
*parvidens*
sp.1
E) PCA of *Parascaptor* species - ventral view of skull
Principal Component 1
Principal Component 2
*leucura*
sp.1
sp.2
F) PCA of *Euroscaptor* species - ventral view of skull
Principal Component 1
Principal Component 2
*grandis*
*klossi*
*kuznetsovi*
*longirostris*
*malayana*
*micrura*
*orlovi*
*parvidens*
sp.1
G) CVA of *Parascaptor* species - ventral view of skull
Canonical Variate 1
Canonical Variate 2
*leucura*
sp.1
sp.2
H) CVA of *Euroscaptor* species - ventral view of skull
Canonical Variate 1
Canonical Variate 2
*grandis*
*klossi*
*malayana*
*micrura*
sp.1

Figure S5. Results of principle component analysis (PCA) and canonical variate analysis (CVA) based on measurement-based morphometric and outline-based geometric morphometric analyses. A-D: PCA plots showing scores on PC1 and PC2 derived from 15 $log_{10}$-transformed craniomandibular variables for (A) *Parascaptor* and (B) *Euroscaptor*, and corresponding CVA plots displaying scores on CV1 and CV2 for the same data in (C) *Parascaptor* and (D) *Euroscaptor*. E-H: PCA plots illustrating scores on PC1 and PC2 based on skull outline shapes for (E) *Parascaptor* and (F) *Euroscaptor*, and corresponding CVA plots showing scores on CV1 and CV2 for the same data in (G) *Parascaptor* and (H) *Euroscaptor*.

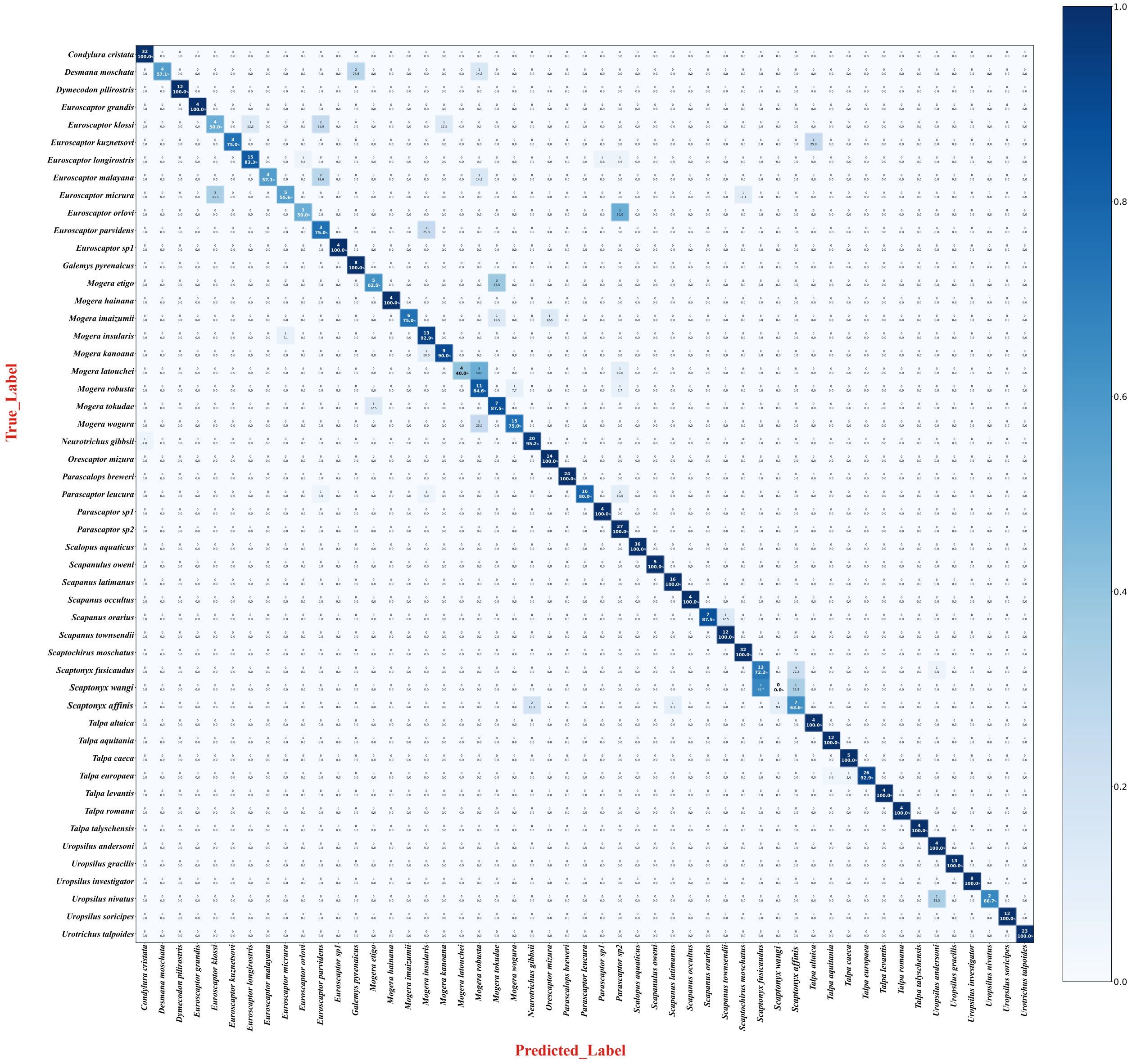

**Figure S6.** Confusion matrix heatmap showing the image-based identification accuracy for recognized and cryptic species across all species and putative species using a one-step strategy. Each photo was directly identified to one of the 51 species. The Y-axis (true_label) represents the actual species classification of each species, while the X-axis (predicted_label) represents the species classification result predicted by the model. Numbers at the top of each cell denote the number of specimens assigned to each species, while the percentage of specimens assigned to that species is listed at the bottom of the cell. The overall accuracy of the model, based on the correct identification of individual photos, was 88.0% (see Table 1).

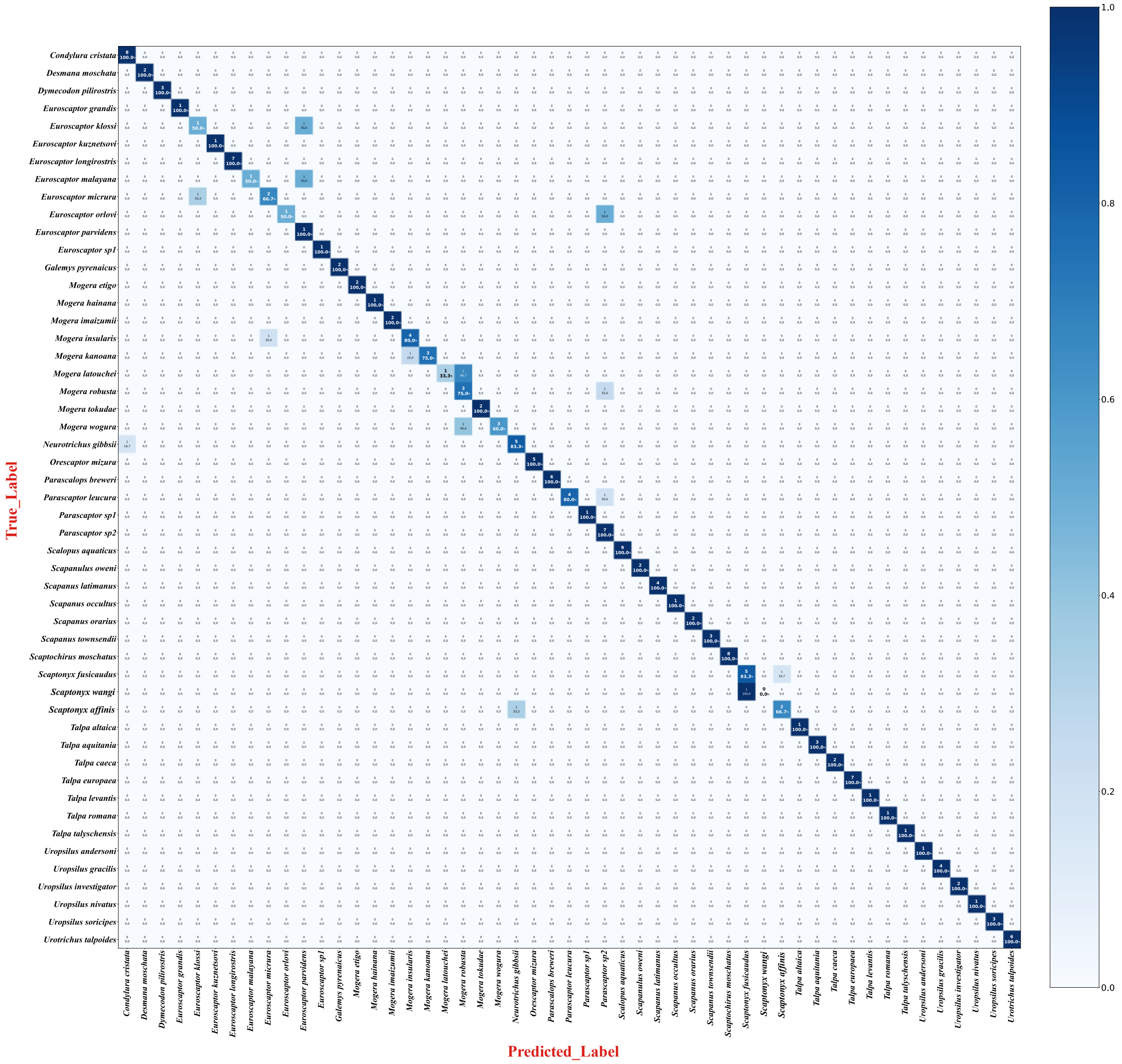


**Figure S7**. Confusion matrix heatmap showing the individual-based identification accuracy for recognized and cryptic species across all species and putative species using a one-step strategy. Each specimen was directly identified to one of the 51 species. The Y-axis (true_label) represents the actual species classification, while the X-axis (predicted_label) represents the species classification result predicted by the model. Numbers at the top of each cell denote the number of specimens assigned to each species, while the percentage of specimens assigned to that species is listed at the bottom of the cell. The overall accuracy of the model, based on the correct identification of individual specimen, was 88.5% (see Table 1).

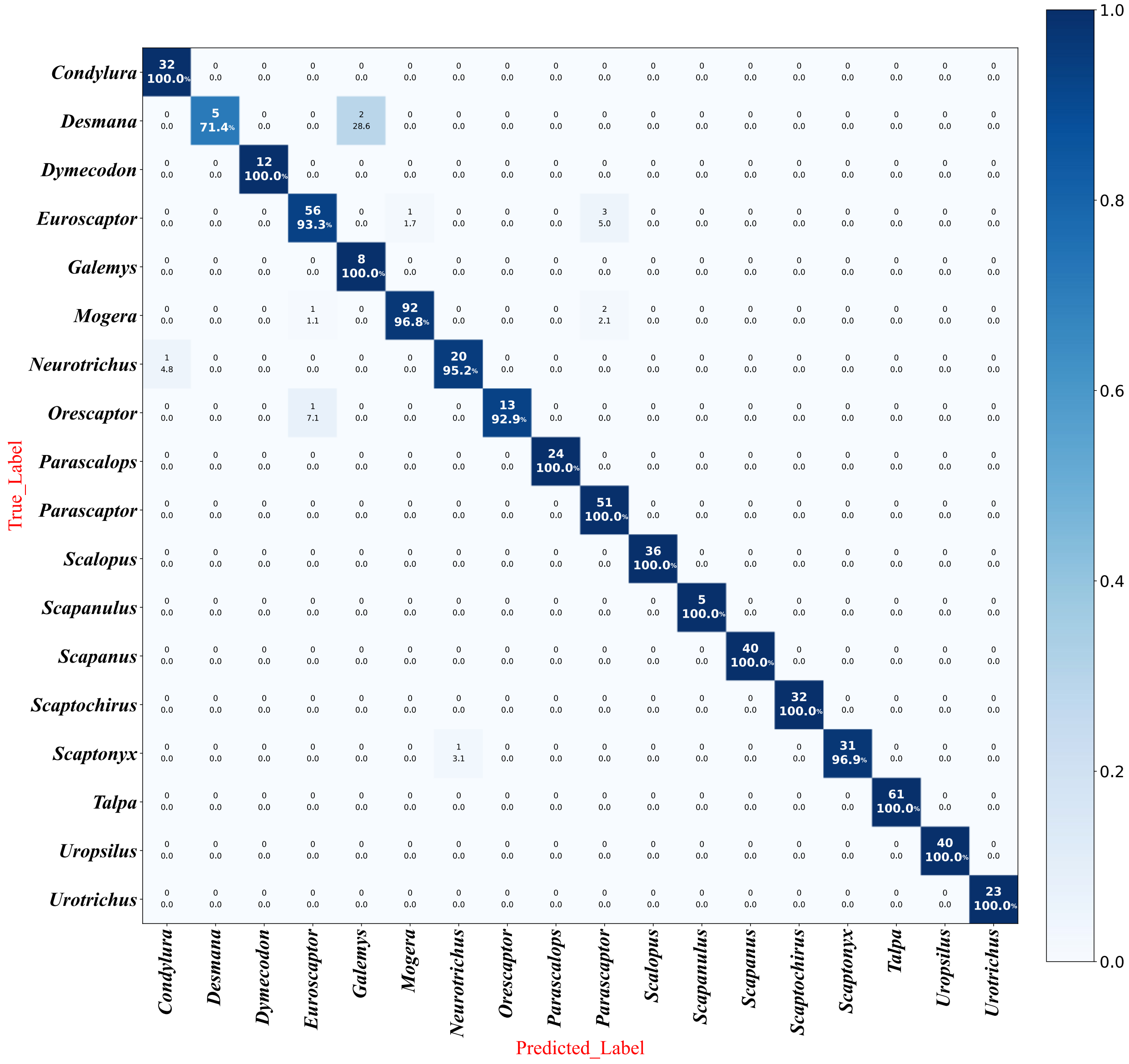


**Figure S8.** Confusion matrix heat map showing the image-based identification accuracy for all genera using HIS-NET. The Y-axis (true_label) represents the actual species classification, while the Y-axis (predicted_label) represents the species classification result predicted by the model. Numbers at the top of each cell denote the number of specimens assigned to each species, while the percentage of specimens assigned to that species is listed at the bottom of the cell. The overall accuracy of the model, based on the correct identification of individual images, was 98.0% (see Table 1).

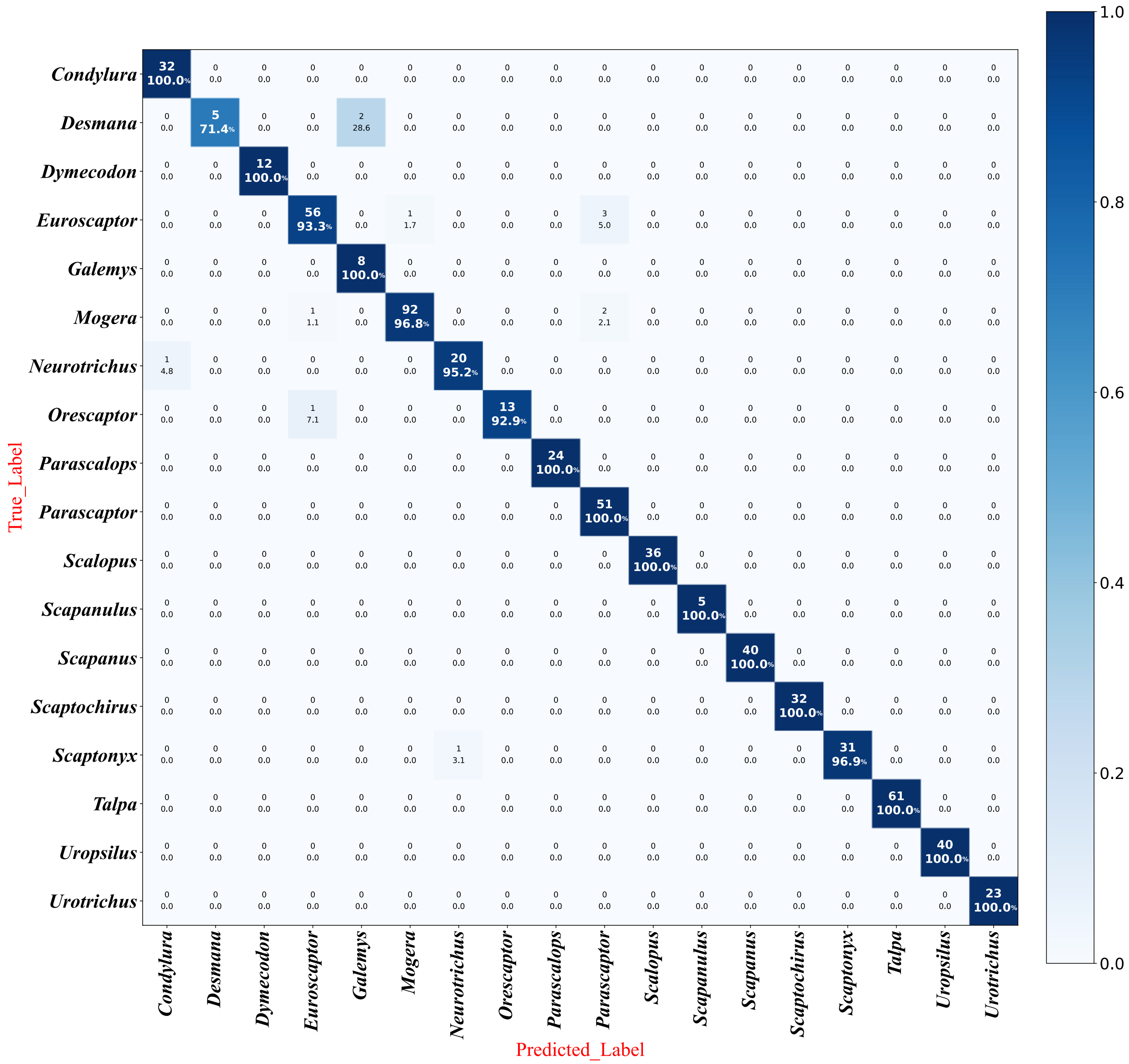


**Figure S9.** Confusion matrix heatmap showing the specimen-based identification accuracy for all genera using HIS-NET. The Y-axis (true_label) represents the actual species classification, while the X-axis (predicted_label) represents the species classification result predicted by the model. Numbers at the top of each cell denote the number of specimens assigned to each species, while the percentage of specimens assigned to that species is listed at the bottom of the cell. The overall accuracy of the model, based on the correct identification of individual specimens, was 97.0% (see Table 1).

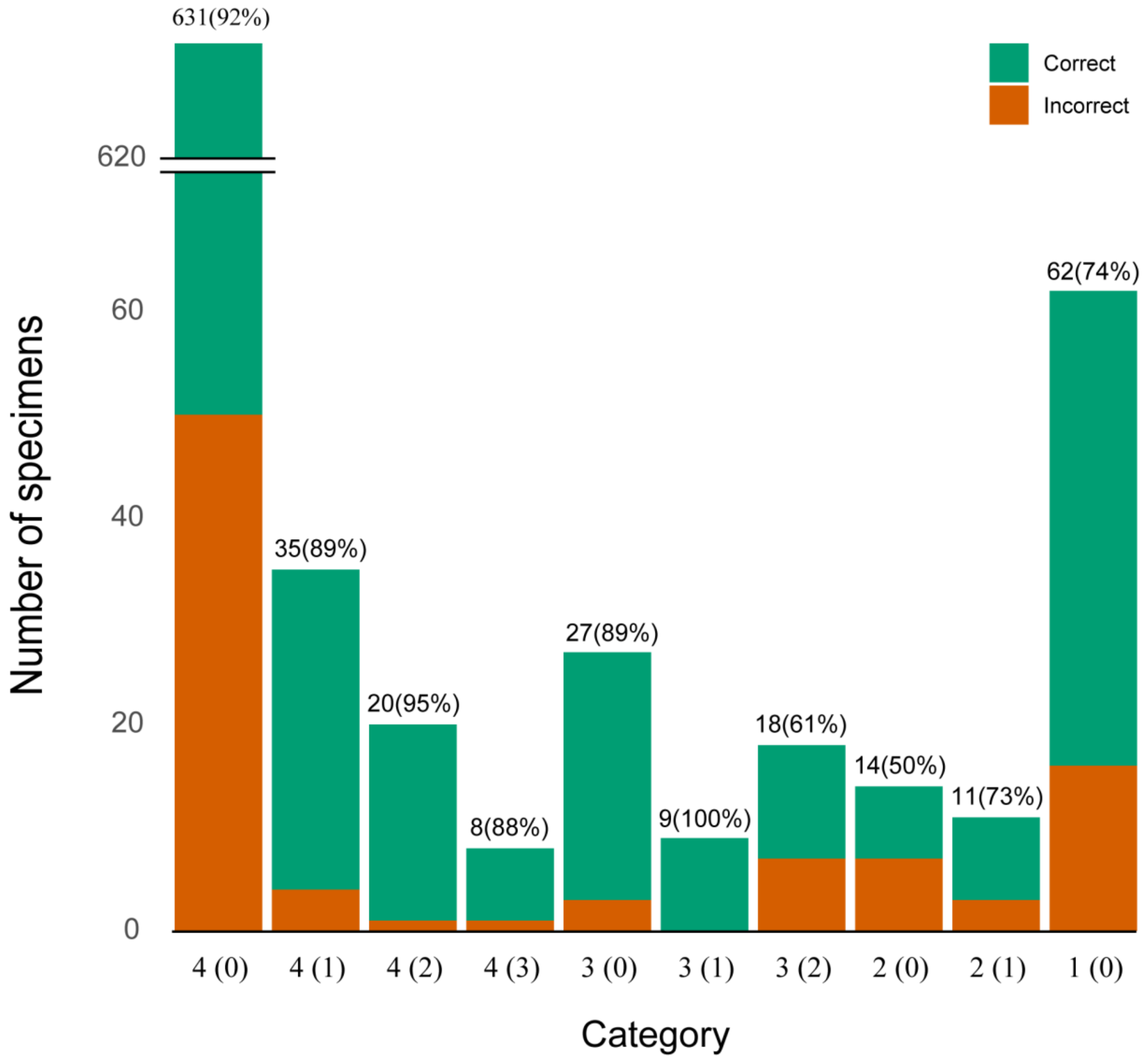


**Figure S10.** Summary of species identification accuracy obtained following five iterations of cross-validation with HIS-NET. Each bar corresponds to a specific category, with values below each bar indicating the number of available images per specimen, and those in parentheses representing the number of images for each specimen showing a broken skull or mandible element. The height of each bar reflects the total number of specimens per category, with the green and orange segments corresponding to the number of correctly and incorrectly identified specimens, respectively. Numbers above each bar indicate the total number of specimens examined per category with the overall percent species identification accuracy for each category given in parentheses.

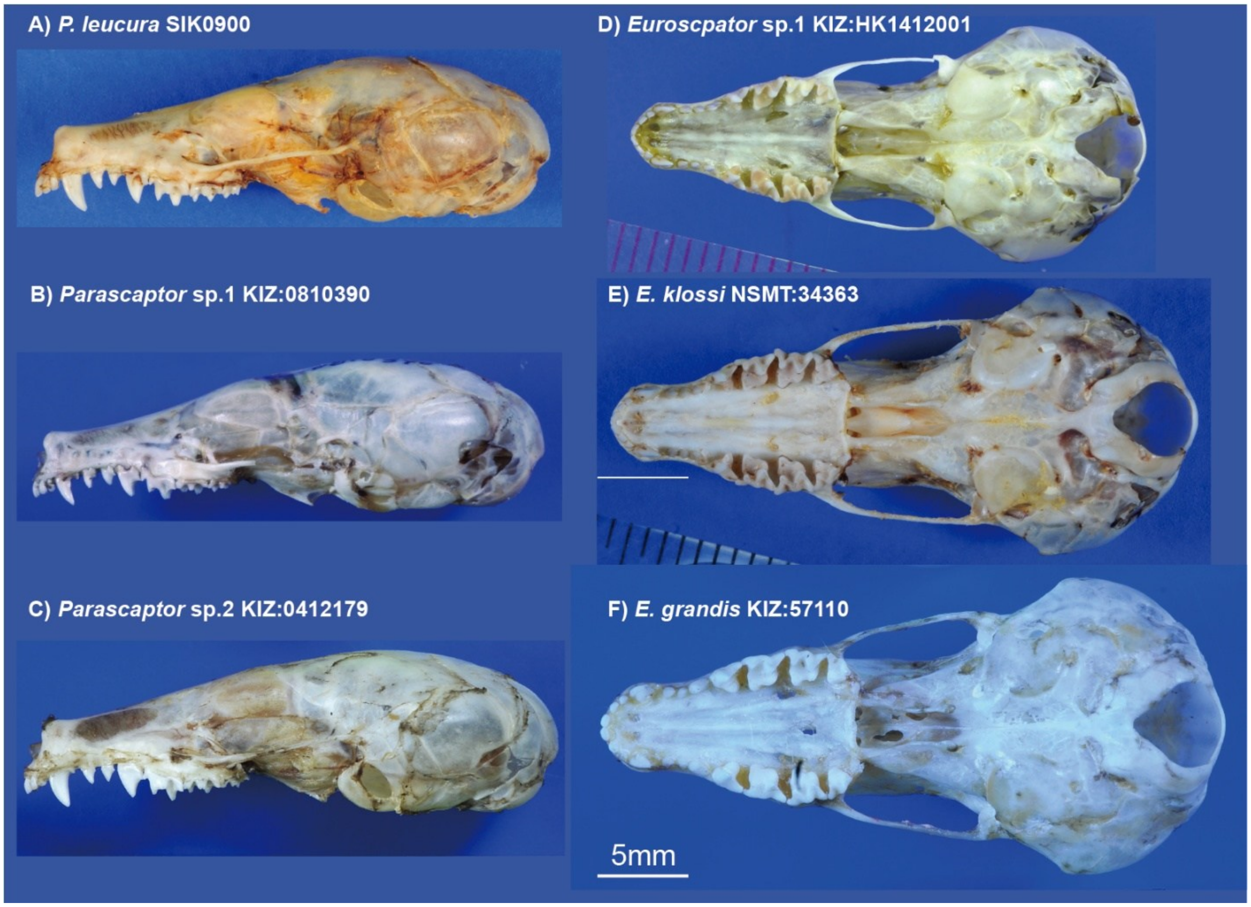


**Figure. S11.** The lateral skull view of (A-C) three *Parascaptor* species/putative species and (D-F) the ventral skull view of three *Euroscaptor* species/putative species.

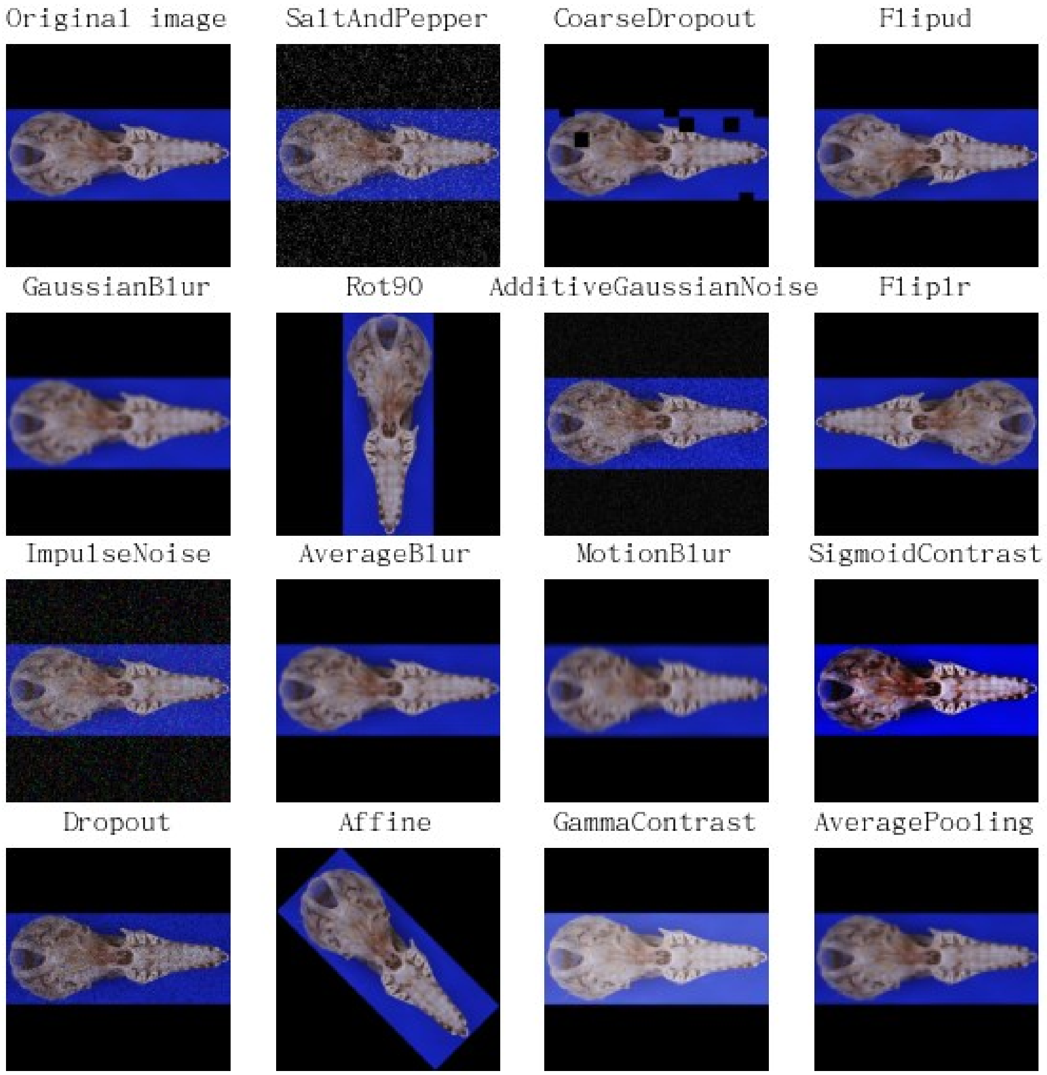


**Figure S12**. Illustration of the various augmentation strategies used in this study. The original image (top left) was subjected to different augmentation techniques to introduce minor distortions and mitigate overfitting during neural network training. The augmentation methods included noise injection (SaltAndPepper, AdditiveGaussianNoise, ImpulseNoise), blurring (GaussianBlur, AverageBlur, MotionBlur), rotation (Rot90, Affine (rotate 45 degrees)), mirroring (Flipud (vertical flip), Fliplr (horizontal flip)), masking (CoarseDropout, Dropout), and contrast adjustment (SigmoidContrast, GammaContrast and AveragePooling).
